# Comparative Analysis of Embryo Proper and Suspensor Transcriptomes in Plant Embryos With Different Morphologies

**DOI:** 10.1101/2020.11.30.404376

**Authors:** Min Chen, Jer-Young Lin, Xiaomeng Wu, Nestor R. Apuya, Kelli F. Henry, Brandon H. Le, Anhthu Q. Bui, Julie M. Pelletier, Shawn Cokus, Matteo Pellegrini, John J. Harada, Robert B. Goldberg

## Abstract

An important question is what genes govern the differentiation of plant embryos into suspensor and embryo-proper regions following fertilization and division of the zygote. We compared embryo proper and suspensor transcriptomes of four plants that vary in embryo morphology within the suspensor region. We determined that genes encoding enzymes in several metabolic pathways leading to the formation of hormones, such as gibberellic acid, and other metabolites are up-regulated in giant Scarlet Runner Bean and Common Bean suspensors. Genes involved in transport and Golgi body organization are up-regulated within the suspensors of these plants as well – strengthening the view that giant specialized suspensors serve as a hormone factory and a conduit for transferring substances to the developing embryo proper. By contrast, genes controlling transcriptional regulation, development, and cell division are up-regulated primarily within the embryo proper. Transcriptomes from less specialized soybean and *Arabidopsis* suspensors demonstrated that fewer genes encoding metabolic enzymes and hormones are up-regulated. Genes active in the embryo proper, however, are functionally similar to those active in Scarlet Runner Bean and Common Bean embryo proper regions. We uncovered a set of suspensor- and embryo-proper-specific transcription factors (TFs) that are shared by all embryos irrespective of morphology, suggesting that they are involved in early differentiation processes common to all plants. ChIP-Seq experiments with Scarlet Runner Bean and soybean WOX9, an up-regulated suspensor TF, gained entry into a regulatory network important for suspensor development irrespective of morphology.

**Significance:** How plant embryos are differentiated into embryo proper and suspensor regions following fertilization is a major unanswered question. The suspensor is unique because it can vary in morphology in different plant species. We hypothesized that regulatory genes controlling the specification of embryo proper and suspensor regions should be shared by all plants irrespective of embryo morphology. We compared embryo proper and suspensor transcriptomes of plants with distinct suspensor morphologies. Scarlet Runner Bean and Common Bean have highly specialized giant suspensor regions, whereas soybean and *Arabidopsis* suspensors are smaller and less specialized. We uncovered a small set of embryo-proper- and suspensor-specific transcription factors shared by all embryos irrespective of morphology, suggesting that they play an important role in early embryo differentiation.

## Introduction

One of the major unsolved questions in plant biology is how regulatory networks embedded in the genome choreograph processes leading to the specification and differentiation of different embryonic regions following zygote formation. In most higher plants, the zygote divides asymmetrically giving rise to an embryo consisting of two regions with distinct developmental fates – the embryo proper and the suspensor (1–3) (Fig. 1). The embryo proper differentiates into regions that enable the next plant generation to develop following seed germination. These include shoot and root meristems that generate the plant body, and cotyledons which serve as an energy source until the germinating seedling is able to survive on its own via photosynthesis (1). By contrast, the suspensor is a terminally differentiated embryonic region that anchors the embryo proper to surrounding seed tissue and degenerates by the time embryogenesis is complete (4–7). One hundred and forty years ago it was known that the suspensor varies greatly morphologically among different plant species, in contrast with the less variant embryo proper (8). Enlarged highly specialized suspensors were speculated to produce substances required for early embryo development (8, 9) – a hypothesis that has stood the test of time (4–7). A series of elegant experiments have illuminated the signaling pathways and regulators responsible for establishing zygotic polarity and directing the embryo to follow embryo proper and suspensor differentiation pathways (3, 10–12). However, most of the regulatory genes and genomic wiring that control these processes are largely unknown.

**Fig. 1.**
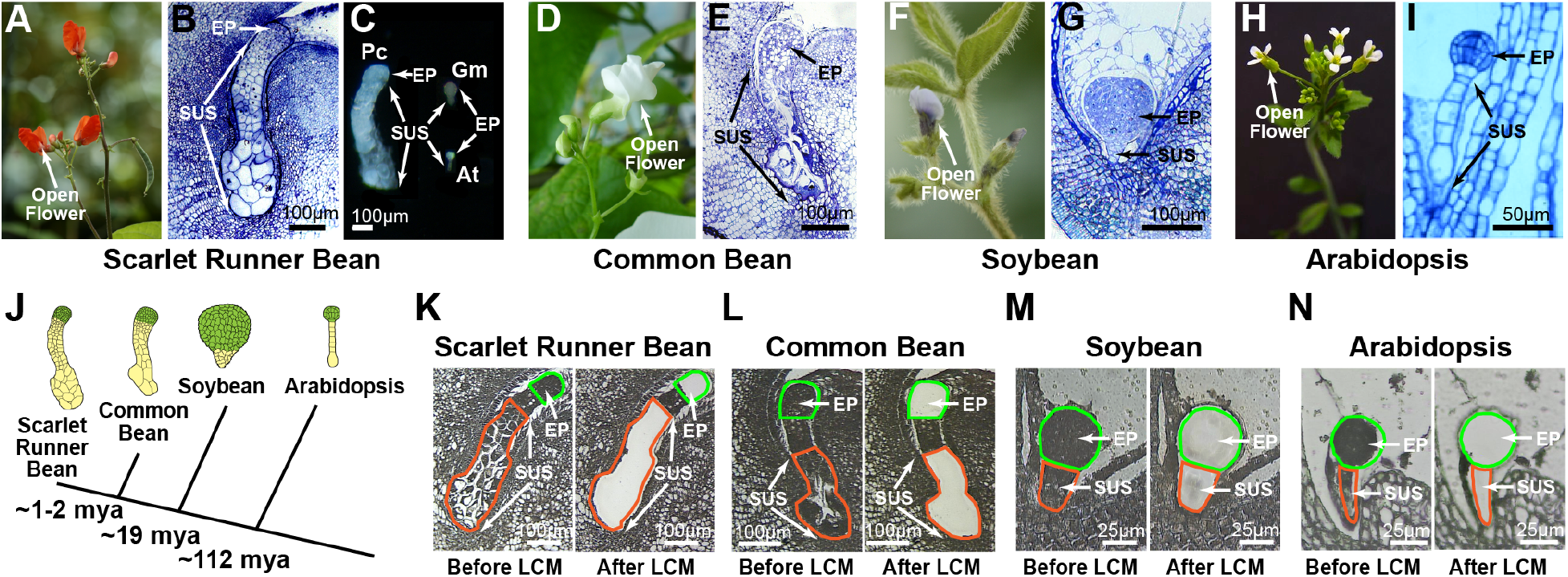
Plant species and embryos used for suspensor and embryo proper transcriptome experiments. *(A)* SRB open flower. *(B)* Plastic section (1 μm) of SRB globular stage embryo. (*C*) Hand-dissected SRB (Pc), *Arabidopsis* (At), and SB (Gm) globular stage embryos. *(D)* CB open flower. *(E)* Paraffin section (6 μm) of CB globular stage embryo. *(F)* SB open flower. *(G)* Paraffin section (6 μm) of SB globular stage embryo. *(H) Arabidopsis* open flower. *(I)* Paraffin section (6 μm) of *Arabidopsis* globular stage embryo. *(J)* Evolutionary relationships between SRB, CB, SB, and *Arabidopsis* (mya, million years ago) (16, 67, 68). Embryo cartoons are not drawn to scale and were traced from paraffin sections of globular stage embryos. Green and yellow colors indicate the embryo proper and suspensor, respectively. *(KN)* Paraffin sections (6 μm) of SRB (*K*), CB *(L),* SB *(M)* and *Arabidopsis (N)* globular stage embryos before and after capture by LCM. EP, embryo proper; SUS, suspensor.

Legume embryos exhibit a wide spectrum of suspensor sizes and shapes (8, 13). For example, Scarlet Runner Bean (SRB) *(Phaseolus coccineus)* and the Common Bean (CB) *(Phaseolus vulgaris*) have large multicellular suspensors (Fig. *1A-E*) that contain polytene chromosomes resembling those of *Drosophila* salivary glands suggesting high metabolic activity (14, 15). These closely related legumes are separated by only 1-2 mya (Fig. 1*J*), and have similar genome sizes and chromosome numbers (16–19). By contrast, soybean (SB) *(Glycine max),* a more distant legume (Fig. 1*J*), has smaller and less specialized suspensors (Fig. *1F* and *G*) resembling those of *Arabidopsis (Arabidopsis thaliana)* (Fig. *1H* and *I*). The large size of SRB and CB embryos permits hand dissection of embryo proper and suspensor regions for experimental studies *(Appendix SI* Fig. S1*A*), which is not practical in SB and *Arabidopsis* (20). Almost five decades ago Ian Sussex, Mary Clutter, Ed Yeung, and their colleagues took advantage of this property to show that giant SRB and CB suspensors are highly active and supply the embryo proper with substances responsible for embryo proper growth (21–23) – sustaining the ideas of pioneering plant embryologists of the late 19th and early 20th centuries (8, 9). Subsequently, experiments with isolated SRB and CB suspensors showed that they contain several hormones, including gibberellic acid (GA), auxin, cytokinin (CK), and abscisic acid (ABA) (24–27), suggesting that hormone signaling might play a critical role in suspensor differentiation and function (24).

We have been using giant SRB embryos as a system to dissect the genomic processes that control suspensor and embryo proper differentiation (28–31). We identified genes that are up-regulated in the suspensor shortly after fertilization – several of which encode enzymes in the GA biosynthetic pathway (28, 30). We dissected the regulatory regions of two suspensor-specific genes – *G564,* encoding a protein of unknown function, and *GA20-oxidase (GA20ox),* specifying an enzyme in the GA biosynthetic pathway (28, 31). Our experiments uncovered a *cis* regulatory module containing five *cis*-elements that are each required for suspensor-specific transcription of the *G564* and *GA20ox* genes. Studies with transgenic plants showed that this regulatory module works in the suspensors of distantly related tobacco and *Arabidopsis* embryos (6), suggesting a conserved suspensor-specific regulatory pathway across the plant kingdom.

Here, we take advantage of the morphological differences between several plant embryos and use laser capture microdissection (LCM) and RNA-Seq to characterize SRB, CB, SB, and *Arabidopsis* embryo proper and suspensor transcriptomes (Fig. 1). Our hypothesis is that irrespective of embryo morphology there is a shared set of embryo-proper and suspensor transcription factors (TFs) that drive the differentiation of these regions and are conserved in higher plants. Our experiments uncovered (i) the spectrum of functional differences between the suspensor and embryo proper in each plant species, (ii) a high degree of metabolic specialization in large SRB and CB suspensors, and (iii) a small set of embryo-proper- and suspensor-specific TFs common to all plant species investigated that might play a major role in early embryo differentiation. Finally, SRB and SB ChIP-Seq experiments with one of the shared suspensor-specific TF mRNAs, WUSCHEL-RELATED HOMEOBOX 9 (WOX9), uncovered several TFs that are putative WOX9 targets, some of which are conserved between both plants. How embryo region-specific TFs are integrated into circuitry required for early embryo differentiation and function remain to be determined.

## Results

### CB Can Be Used as an SRB Reference Genome

We examined whether we could use CB as a reference genome for SRB expression data because these two legumes are separated by only 1-2 mya (Fig. 1*J*), and an excellent CB genome draft exists (32). We generated several hundred thousand SRB Expressed Sequence Tags (ESTs) from hand dissected globular stage embryo proper and suspensor regions *(SI Appendix* Fig. S1*A*), and aligned these ESTs with the predicted transcripts of CB genes. Approximately 15,000 diverse transcripts were represented in our embryo EST population, and there was >95% similarity between SRB and CB sequences (*SI Appendix* Fig. SI*B* and *C*), reflecting the close evolutionary relationship between SRB and CB at the gene level.

We used LCM to isolate embryo proper and suspensor regions from SRB globular stage embryos (Fig. *1K* and *SI Appendix* Fig. S1*D*), and uncovered the spectrum of transcripts present in each of these embryonic regions using RNA-Seq. Approximately 95% of our EST collection was represented in the RNA-Seq reads (*SI Appendix* Fig. S1*E*). To ensure that the CB genome can be used as a reference for SRB expression data, we sequenced the SRB genome at approximately 100X coverage (*SI Appendix* Fig. S1*F*), and compared the alignments of RNA-Seq reads with both CB and SRB genome sequences (*SI Appendix* Fig. S1*G*). The same number of mappable reads were obtained with both genomes (*SI Appendix* Fig. S1*G*), indicating that the CB draft genome and annotated genes can be used as a reference for analyzing SRB RNA-Seq data.

### Absence of Contamination From Surrounding Seed Tissue in Embryo Proper and Suspensor Regions Captured By LCM

We used SRB embryo medial sections to avoid potential contamination from surrounding seed tissue during embryo proper and suspensor region LCM (Fig. 1*K*) (see Materials and Methods) (20, 33). We searched our embryo proper and suspensor RNA-Seq reads for endothelium-specific transcripts (G563) (*SI Appendix* Fig. S2*A*), seed coat-specific transcripts (GA2-oxidase) (*SI Appendix* Fig. S2*B*), and endosperm-specific transcripts (AGL62) that we (G563 and GA2-oxidase) (28, 30), and others (AGL62) (34), identified previously. We compared the RNA-Seq reads for these transcripts with those of ent-kaurene oxidase (KO), a suspensor-specific mRNA (Fig 2 and *SI Appendix* Fig. S2*C*) (28). There were few, if any, RNA-Seq reads for surrounding seed tissue transcripts in comparison with the KO control (*SI Appendix* Fig. S2*D* and *E*), indicating that our embryo proper and suspensor LCM sections had little, or no, contaminating transcripts from adjacent seed tissues.

**Fig. 2.**
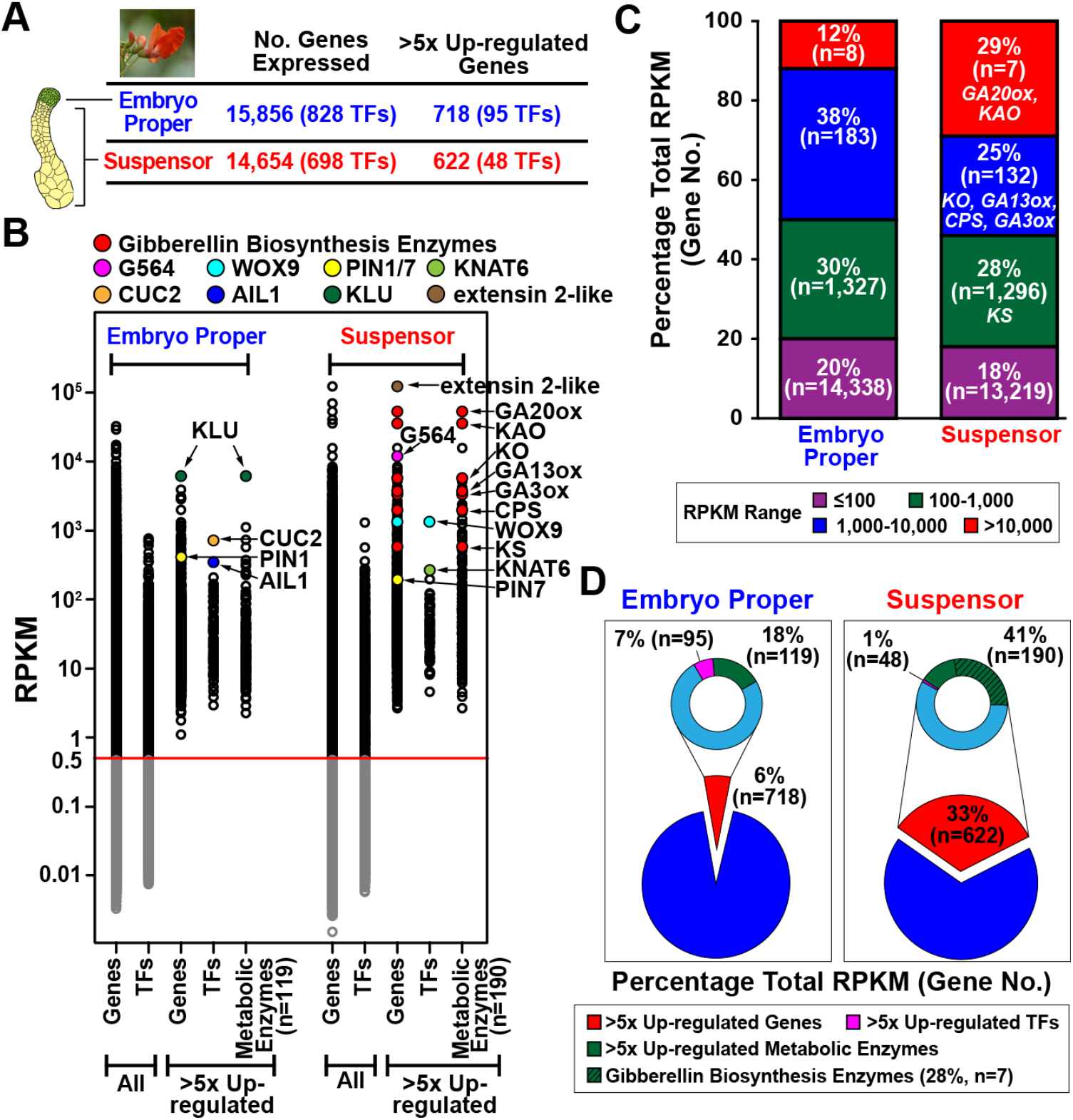
Gene activity in SRB suspensor and embryo proper regions. *(A)* Genes active (>0.5 RPKM) in at least one biological replicate, and >5 fold up-regulated genes in embryo proper and suspensor (see Materials and Methods). All expressed genes and those up-regulated >5-fold are listed in Dataset S1. TFs, transcription factor mRNAs. *(B)* The abundance distribution of mRNAs in different embryo proper and suspensor mRNA populations represented by Reads Per Kilobase of Transcript per Million Reads Mapped (RPKMs) (35). Each circle represents a different mRNA sequence. RNAs with <0.5 RPKM in both biological replicates were not used in subsequent analyses, as indicated by the grey circles below the red line. Enzyme mRNAs were identified using the Plant Metabolic Pathway Database (https://www.plantcyc.org). *(C)* Percentage of embryo proper and suspensor mRNA populations with different prevalence levels. Parentheses indicate the number of different mRNAs in each prevalence group. *(D)* Percentage of total RPKM for different embryo proper and suspensor mRNA populations. Parentheses represent the number of mRNAs in each population. Gene abbreviations are defined in *SI Appendix* Table S1.

### SRB Embryo Proper and Suspensor Transcriptomes Contain a Spectrum of mRNAs

Approximately 15,000 diverse mRNAs were present in each embryo region, including 700-800 TF mRNAs (Fig. 2*A* and Dataset S1). The union of these mRNA sets indicated that there were 17,500 diverse mRNAs in the embryo as a whole – including 1,000 TF mRNAs and a small set of transcripts specific for embryo proper and suspensor regions. We filtered the RNA-Seq reads to include only those with RPKM values >0.5 (Fig *2B*). We estimated that this criterion scored mRNAs at functionally meaningful mRNA levels of >1 molecule/cell (35, 36). Both the embryo proper and suspensor mRNA populations spanned a wide range of prevalences consistent with those found in plant embryos (37). TF mRNA prevalences also spanned several orders of magnitude, suggesting that a range of regulatory inputs are required by each embryo region (Fig. 2*B*).

A small fraction of both the embryo proper and suspensor mRNA mass (~20%) contained most of the diverse mRNAs (Fig. *2C*). On average, these transcripts were represented only a few times per cell (~15) and resembled complex class rare mRNAs typical of plant embryo cells (37). The suspensor, however, contained a large fraction of highly prevalent mRNAs present in tens of thousands of copies per cell as compared with the embryo proper (Fig. 2*B* and *C*). Almost 30% of the suspensor mRNA mass contained only seven different sequences, including those encoding the GA biosynthesis enzymes GA20ox and KAO (Fig. 2*C*). In fact, GA20ox and KAO mRNAs were among the most prevalent mRNAs in the entire embryo (Fig. 2*B* and *C*), suggesting a high degree of metabolic specialization within the suspensor.

### SRB Embryo Proper and Suspensor Regions Contain Specific mRNA Sets

We used EdgeR (FDR <0.05) to identify mRNAs that were >5-fold more prevalent, or up-regulated, in each embryo region (see Materials and Methods). We uncovered 718 and 622 embryo-proper- and suspensor-specific mRNAs, respectively (Fig. 2*A* and Dataset S1), and then identified mRNAs encoding metabolic enzymes and TFs in each up-regulated mRNA set (Fig. *2B* and *D*, and Dataset S1). Region-specific mRNAs spanned a range of prevalences analogous to those in the unselected populations (Fig. 2*B*). Suspensor-specific mRNAs, however, occupied a greater proportion of mRNA mass than their embryo proper counterparts (33% vs. 6%), reflecting the presence of highly prevalent mRNAs (Fig. 2*B-D*). Over 40% of the suspensor-specific mRNA mass consisted of 190 diverse metabolic enzyme mRNAs, including those encoding GA biosynthesis enzymes (Fig. 2*D*). By contrast, only 18% of the embryo-proper-specific mRNA mass encoded 119 metabolic enzymes (Fig. 2*D*), indicating that a greater proportion of gene activity within the suspensor is directed towards specialized metabolic processes.

In contrast with mRNAs involved in metabolism (Fig. 2*D*), TF mRNAs represented a larger fraction of the embryo-proper up-regulated mRNA set compared with the suspensor (7% vs. 1%) (Fig. 2*D*). We uncovered 95 and 48 diverse TF mRNAs specific to the embryo proper and the suspensor, respectively (Fig. 2*A* and Dataset S1), representatives of which are listed in Fig. 3. The precise roles that the majority of these region-specific TF mRNAs perform within the SRB embryo proper and suspensor at the globular stage are unknown. However, they include TF mRNAs with known developmental and hormone signaling functions (Fig. 3*A* and Dataset S1). For example, embryo-proper-specific TF mRNAs include those involved in shoot meristem development [SHOOT MERISTEMLESS (STM)] and cotyledon separation [CUP-SHAPED COTYLEDON 2 (CUC2)] (12), among others (Fig. 3*B* and Dataset S1). By contrast, the suspensor-specific TF mRNA set contains WOX9, AUXIN RESPONSE FACTOR 16 (ARF16), and HOMEODOMAIN GLABROUS 11 (HDG11) TFs that are required for suspensor specification (Fig. 3*C* and Dataset S1) (3). The most prevalent up-regulated TF mRNAs in the embryo proper and suspensor were CUC2 and WOX9, respectively (Fig. 2*B*). A higher percentage (80%) of embryo-proper-specific TFs play a role in developmental and hormone response processes compared with their counterparts in the suspensor (40%) (Fig. 3*A* and *SI Appendix* Fig. S3), indicating significant functional differences between the embryo-proper- and suspensor-specific TF mRNA populations.

**Fig. 3.**
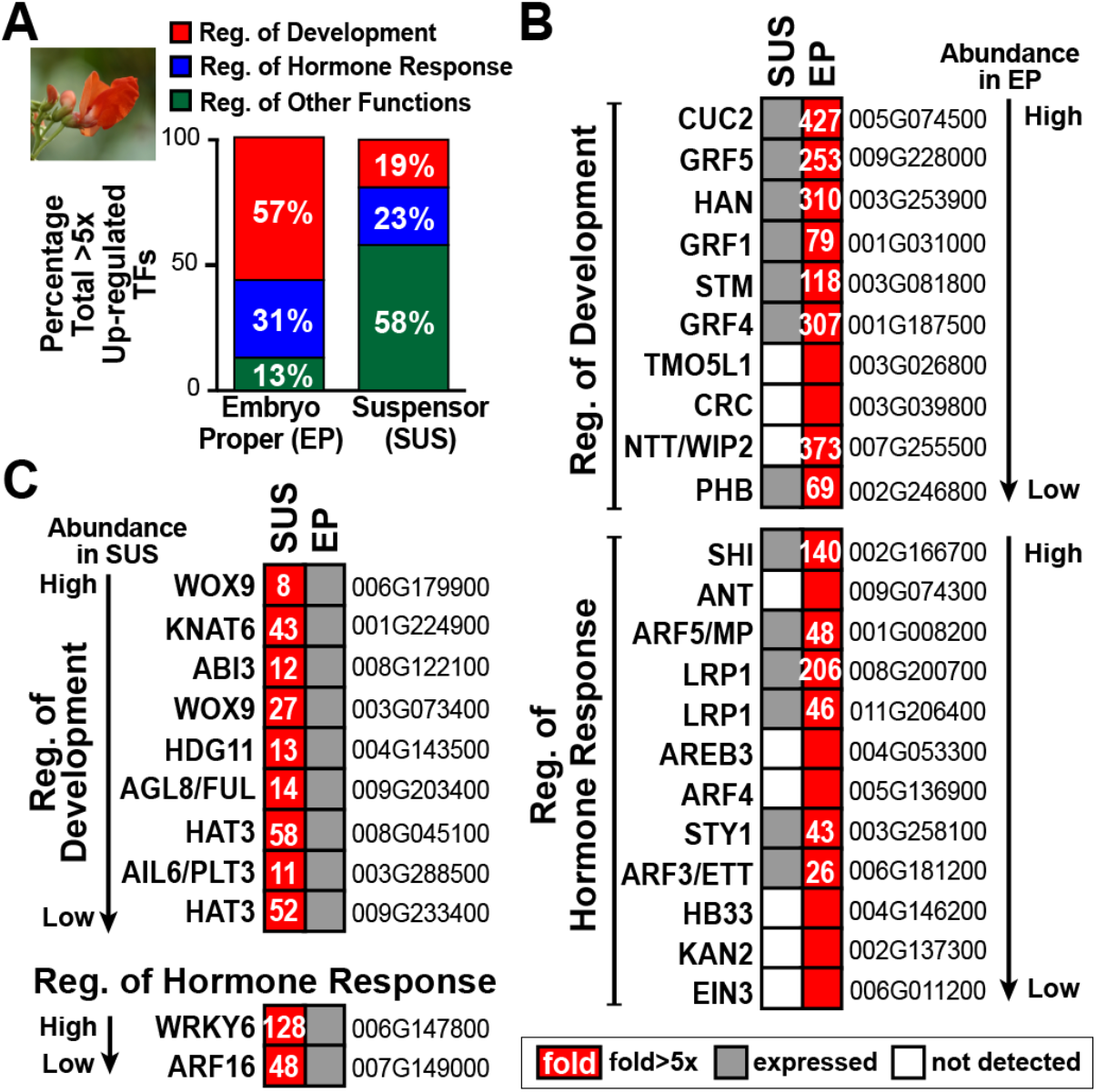
Representative SRB up-regulated suspensor and embryo proper TF mRNAs. *(A)* Percentage of TF mRNAs in different functional groups (Fig. 4 and *SI Appendix* Fig. S3). *(B* and *C)* Representative up-regulated embryo proper (*B*) and suspensor (*C*) TF mRNAs listed by prevalence. Numbers in red squares represent fold differences. Red squares without a number indicate that mRNAs were not detected in the other embryo region (white squares). Grey squares indicate mRNAs were detected at levels >0.5 RPKM. Gene identifiers (i.e., Pv number) given to right. Gene abbreviations are defined in *SI Appendix* Table S1.

### SRB Embryo Proper and Suspensor Regions Differ Significantly in Biological Processes

We performed Gene Ontology (GO) analysis (FDR <0.05) on up-regulated embryo-proper- and suspensor-specific mRNAs to characterize the major functions that are carried out in each embryo region (Fig. 4). A summary of up-regulated mRNAs in major GO categories is presented in *SI Appendix* Fig. S3, and Dataset S2 contains a list of all embryo proper and suspensor GO terms.

**Fig. 4.**
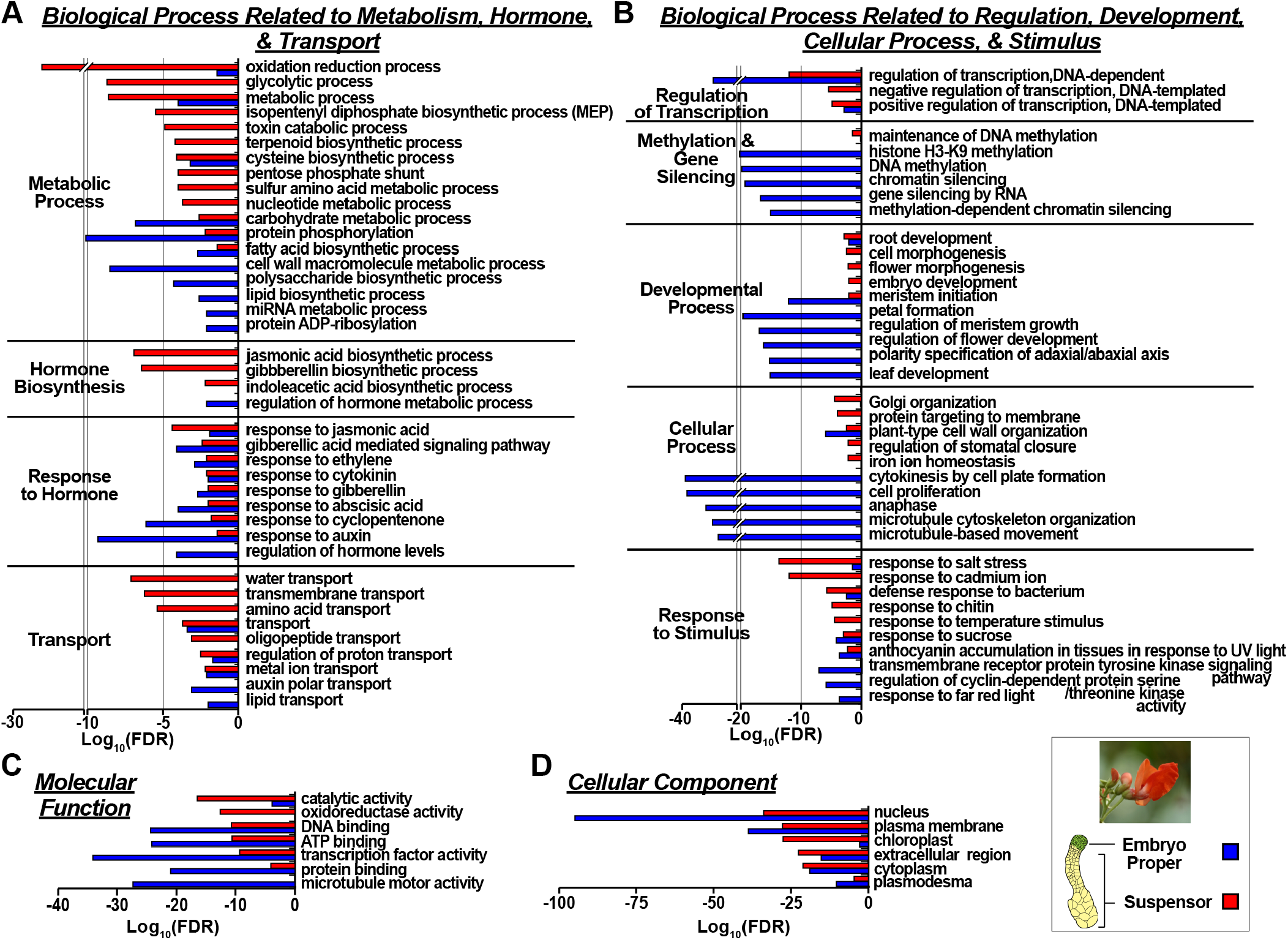
Gene Ontology (GO) terms that are enriched in SRB globular stage embryo proper and suspensor up-regulated mRNAs. (*A* and *B*) Enriched biological process GO terms related to metabolism, hormone, and transport (*A*) and regulation, response to regulation, development, cellular process, and stimulus (*B*). (*C* and *D*) Enriched molecular function (*C*) and cellular component (*D*) GO terms. Only the top five to 10 GO terms are listed. All GO terms are listed in an interactive format in Dataset S2. FDR, false discovery rate.

#### Embryo Proper

The most significant GO terms reflected the cell division, differentiation, and developmental regulatory processes occurring in this embryo region during the globular stage. These included regulation of transcription, transcription factor activity, histone and DNA methylation, cytokinesis by cell plate formation, and cell proliferation, among others (Fig. 4*A*-*C*). Developmental GO terms, such as meristem initiation and polarity specification of the adaxial/abaxial axis reflected patterning events taking place within the embryo proper region. The most significant metabolic process GO terms were carbohydrate, lipid, and cell wall macromolecule (polysaccharide) biosynthesis, although less so than regulatory events. Finally, signaling pathways were prominent among embryo proper GO terms, including polar transport and response to a spectrum of hormones, such as GA, JA, CK, ABA, and auxin. The latter hormone being transported within the embryo proper in a basal direction towards the suspensor by the PIN FORMED 1 (PIN1) transporter to form an auxin gradient that triggers root pole differentiation within the embryo proper (Fig. 2*B* and *SI Appendix* Fig. S4) (38). GO terms for other transporters, including those for lipids and metal ions, were also overrepresented in the embryo-proper-specific mRNA set (Fig. 4*A*). Specific embryo proper mRNAs encoding these transporters, and others (e.g., PIN1), are summarized in *SI Appendix* Fig. S4.

#### Suspensor

A striking aspect of the suspensor was the large number of GO terms reflective of processes related to metabolism, hormone synthesis, and transport, in contrast with the embryo proper. For example, oxidation reduction process and catalytic activity were the most significant biological process and molecular function GO terms, respectively (Fig. 4*A* and *C*). By contrast, there were fewer regulatory and developmental GO terms, and cell proliferation GO terms were absent reflecting the cessation of suspensor cell division at the globular stage (22). Major metabolic pathways such as glycolysis, pentose phosphate shunt, and the synthesis of several hormones, including JA, GA, and indoleacetic acid (auxin), were significant GO terms (Fig. 4*A*). GO terms for response to these hormones were also observed (Fig. 4*A*), indicating the presence of hormone signal transduction pathways within the suspensor. The plastid cellular component GO term reflected the location of several major metabolic processes (e.g., GA biosynthesis) within this organelle (Fig. 4*D*). Finally, transmembrane, golgi organization, water, ion, amino acid, and oligopeptide transport were major GO terms (Fig. 4*A* and *B*). These were encoded by many up-regulated suspensor-specific transporter mRNAs distinct from those present in the embryo proper, including PIN FORMED 7 (PIN7) which is involved in auxin transport and suspensor development (Fig. 2*B* and *SI Appendix* Fig. S4) (3).

### SRB Suspensor mRNAs Encoding Enzymes in Several Interconnected Biosynthetic Pathways Leading to Hormone Production are Up-Regulated

#### Methyl Erythritol-4-Phosphate (MEP) Isoprenoid and GA Pathways

mRNAs encoding enzymes in the plastid-localized MEP isoprenoid pathway were up-regulated within the suspensor (Fig. 5*A* and *SI Appendix* Fig. S5*A*). MEP mRNAs, together with those required for GA biosynthesis (Fig. 2*B*, Fig. 5*A*, and *SI Appendix* Fig. S5*B*), indicated that mRNAs encoding enzymes in entire plastid-localized metabolic pathway from pyruvate to bioactive GA1 and GA4 were up-regulated within the suspensor (Fig. 5*A*), and responsible for generating their GO terms (Fig. 4).

**Fig. 5.**
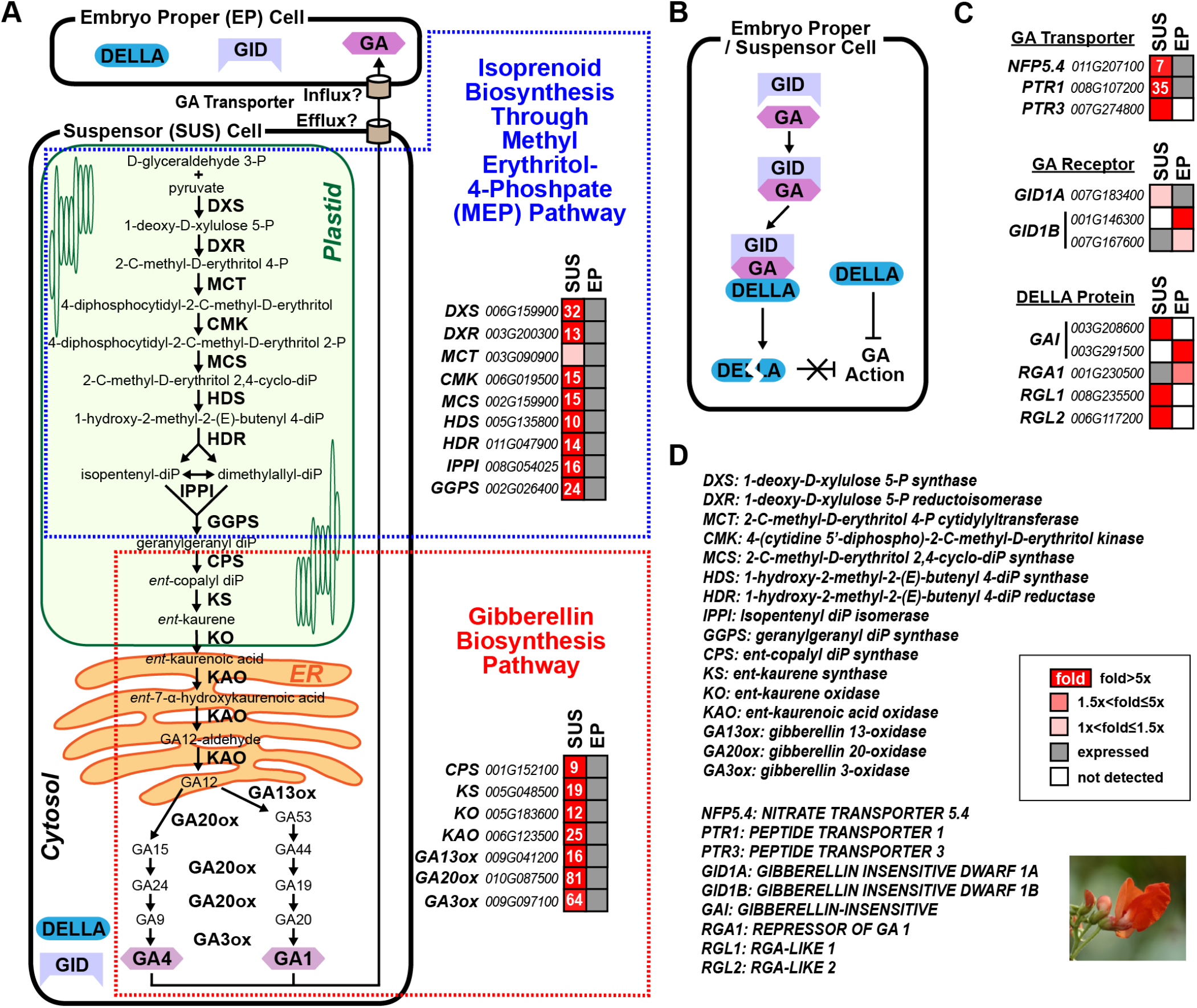
GA biosynthesis and isoprenoid pathway mRNAs in SRB suspensor and embryo proper regions. *(A)* GA and methyl erythritol-4-phosphate (MEP) pathways, and fold-change levels (number in red squares) of pathway mRNAs. GA pathway, MEP pathway, and enzyme intracellular locations were taken from published information (69–72). Enzyme intracellular localizations were confirmed using the DeepLoc machine learning tool (64) (see Materials and Methods and Dataset S3). Suspensor and adjacent embryo proper cell model is based on representation of enzyme, receptor, and transporter mRNAs in these embryo regions (*A* and *C).* GA efflux and influx from the suspensor to the embryo proper is based on classical experiments with SRB embryos suggesting that GA is transferred from the suspensor to embryo proper (41, 73). *(B)* The interaction of GA, GA receptors (GID), and DELLA in GA signaling pathway (74). (*C*) Representation of GA transporter, receptor, and DELLA mRNAs in suspensor and embryo proper. *(D)* GA biosynthesis enzyme, MEP pathway enzyme, and GA signaling protein abbreviations.

We examined mRNAs specifying proteins in the GA signal transduction pathway (Fig. 5*B*). DELLA mRNAs, such as those encoding GIBBERELLIN INSENSITIVE (GAI) and REPRESSOR OF GA-LIKE 1 and 2 (RGL1 and RGL2), were up-regulated in the suspensor (Fig. 5*C*). mRNAs encoding the GA receptor, GIBBERELLIN INSENSITIVE DWARF 1A (GID1A), and GA transporters, NITRATE TRANSPORTER 5.4 (NPF5.4) and PEPTIDE TRANSPORTER 1 and 3 (PTR1 and PTR3), were up-regulated in the suspensor as well (Fig. 5*C*) Significantly, different paralogs of the DELLA GAI and RGA1 mRNAs and GA receptor GID1A and GID1B mRNAs were up-regulated in the embryo proper (Fig. 5*C*). These data suggest that GA is (i) synthesized in the suspensor, (ii) transported to the embryo proper (Fig. 5*A*) and (iii) elicits responses in both embryonic regions (Fig. 4). This is consistent with experiments carried out decades ago that detected the presence of bioactive GA in the SRB suspensor (39), and suggested that GA moves from the suspensor to the embryo proper affecting its development (40, 41).

#### Glycolysis and Pentose Phosphate Shunt Pathways

Plastid-localized glycolysis and the pentose phosphate shunt utilize starch as a substrate and are required to produce precursor molecules for the isoprenoid and GA biosynthetic pathways, and other hormones such as JA, CK, ABA, and auxin (Fig. 6). We examined mRNAs encoding enzymes in glycolytic and pentose phosphate shunt pathways, and found that one or more paralogs of these enzyme mRNAs were either up-regulated or detected within the suspensor (*SI Appendix* Fig. S5*C* and *D*, and Fig. S6). For example, phosphoglucose isomerase (GPI) and glucose-6-phosphate dehydrogenase (G6PD) mRNAs, encoding rate-limiting enzymes of the glycolytic and pentose phosphate shunt pathways, respectively, were up-regulated >5-fold (*SI Appendix* Fig. S5*C* and *D*, and Fig. S6). In addition, mRNAs for the first three enzymes in starch biosynthesis were up-regulated (e.g., ADP glucose pyrophosphorylase large subunit) (Datasets S1 and S2), and starch granules are present in SRB suspensor cells (21). These data suggest a remarkable coordination of metabolic events within the suspensor beginning with starch biosynthesis and culminating in pyruvate, glyceraldehyde-3-phosphate, and chorismite precursors required for several hormone biosynthetic pathways (Fig. 6).

**Fig. 6.**
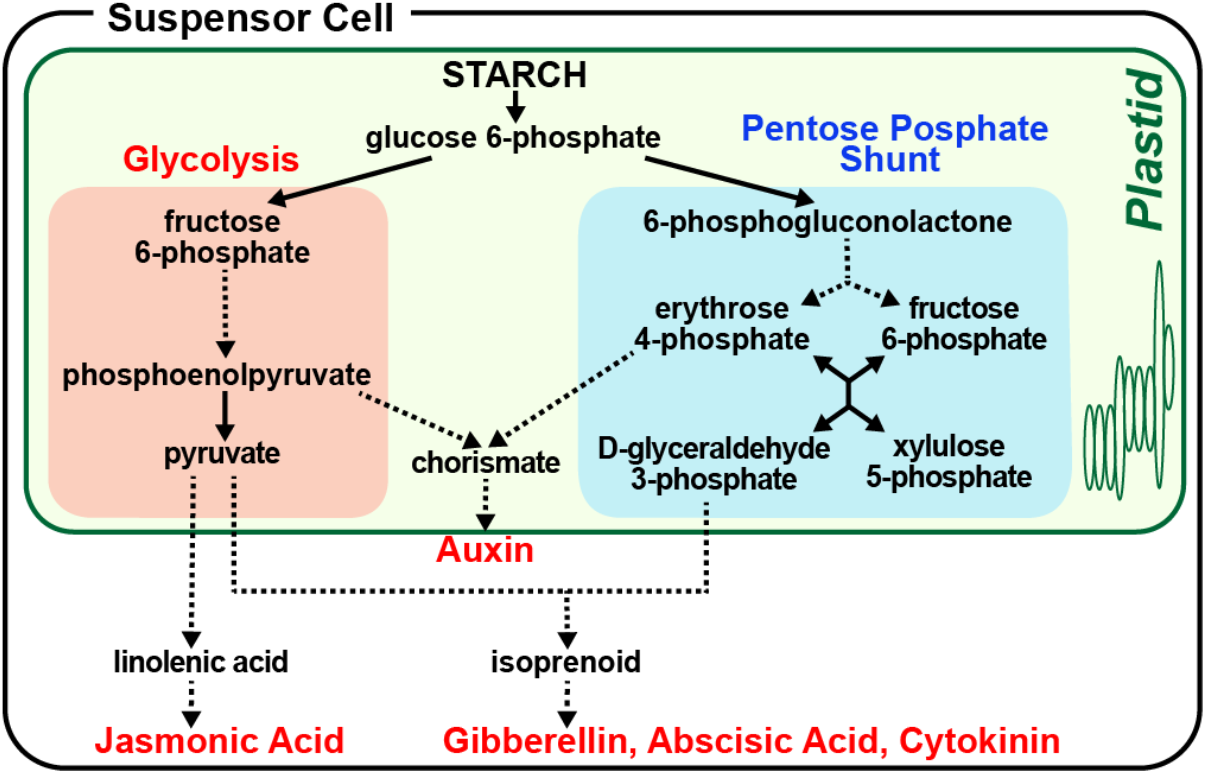
Conceptual overview of glycolysis and pentose phosphate shunt metabolic pathways leading to hormone biosynthesis. Information for pathways taken from references (69, 75).

#### JA, Auxin, ABA, and CK Hormone Pathways

We detected suspensor GO terms for JA and auxin biosynthesis (Fig. 4), suggesting that mRNAs encoding enzymes in these hormone pathways were also up-regulated in the suspensor. Previously, others demonstrated the presence of auxin, ABA, and CK within the SRB suspensor (24, 26, 27), although their sites of synthesis within the embryo were not known. We examined mRNAs encoding enzymes in the JA, auxin, ABA, and CK biosynthetic pathways to determine whether the suspensor had the potential for synthesizing these hormones (*SI Appendix* Fig. S5*E-I*, Fig. S7, and Fig. S8). In each hormone biosynthesis pathway, all of the enzyme mRNAs required for catalytic steps from precursor to final product were detected in the suspensor (*SI Appendix* Fig. *S5E-I*, Fig. S7, and Fig. S8), and many were up-regulated, although not to the levels of isoprenoid and GA biosynthesis mRNAs (*SI Appendix* Fig. S5). We also detected receptor, regulatory, and transporter mRNAs for JA, auxin, ABA, and CK in the suspensor mRNA population (*SI Appendix* Fig. S7 and Fig. S8). For example, several mRNA paralogs of the JA receptor JASMONATE ZIM-DOMAIN PROTEIN (JAZ) were detected, and one, JAZ10, was up-regulated ten-fold compared with the embryo proper (*SI Appendix* Fig. S7). In addition, JA regulator mRNAs, NOVEL INTERACTOR OF JAZ (NINJA) and CORONATINE INSENSITIVE 1 (COI1), were also up-regulated in the suspensor (*SI Appendix* Fig. S7). Similarly, the auxin receptor, TRANSPORT INHIBITOR RESPONSE 1 (TIR1) mRNA, and the auxin response TF mRNA, ARF16, were up-regulated within the suspensor along with the PIN7 efflux carrier mRNA (*SI Appendix* Fig. S4 and Fig. S7). These data, together with hormone response GO terms (Fig. S4), suggest that, in addition to GA, the SRB suspensor has the machinery for synthesizing, transporting, and utilizing JA, ABA, CK, and auxin in signal transduction pathways.

### Gene Expression Activities in the CB Embryo Proper and Suspensor Are Similar to Those in SRB

Developmental and ultrastructure studies by others showed that CB and SRB embryos are indistinguishable from each other, except that the CB embryo is slightly smaller (Fig. 1 *B* and *E*) (42, 43). We used LCM to capture CB globular-stage embryo proper and suspensor regions (Fig. 1*L*), and then profiled their mRNAs using RNA-Seq (Datasets S1 and S2). The embryo proper and suspensor mRNA populations were similar to corresponding SRB mRNAs in every feature, including: (i) similar numbers of genes (16,500) and TFs (900) (*SI Appendix* Fig. S9*A*), (ii) small sets of embryo proper and suspensor specific mRNAs (*SI Appendix,* Fig. S9*A*), (iii) abundance distributions (*SI Appendix* Fig. S9*B*), (iv) presence of highly prevalent suspensor mRNAs, including those encoding GA20ox and KAO (*SI Appendix* Fig. S9*B* and *D*), (v) upregulation of metabolic pathway mRNAs, such as those for MEP, GA biosynthesis, pentose phosphate shunt, and glycolysis (*SI Appendix* Fig. S9*C*, Fig. S11*A*, and Dataset S1), (vi) profile of significant GO terms – including embryo proper enrichment for developmental, transcriptional, and cell proliferation processes; and suspensor enrichment for metabolic and hormone biosynthesis processes (*SI Appendix* Fig. S10 and Dataset S2), (vii) presence of GA signaling proteins in the suspensor (*SI Appendix* Fig. S11*A*), (viii) similar distribution of embryo proper- and suspensor-specific transporters (*SI Appendix* Fig. S11*B-D*), and (ix) similar profiles of embryo-proper- and suspensor-specific TF mRNAs, including the presence of STM and WOX9 mRNAs in the embryo proper and suspensor, respectively, among others (*SI Appendix* Fig. S9*E-G* and Dataset S1). Together, these data indicate that SRB and CB globular stage embryos are virtually indistinguishable in their gene expression profiles and specialized activities – as predicted by their similar morphology and close evolutionary relationship (Fig. 1).

### Gene Expression Activities in SB and *Arabidopsis* Globular Stage Embryos Differ From Those in SRB and CB

SB and *Arabidopsis* suspensor regions are much simpler than those of SRB and CB (Fig. 1). At the globular stage, the *Arabidopsis* suspensor consists of a linear file of 5-7 small cells, and is approximately 200 times smaller than SRB and CB suspensors (Fig. 1*C* and *I*) (44). On the other hand, the SB suspensor consists of a small collection of cells shaped in a V-like structure at the globular stage (Fig. *1G*) that will elongate into a narrow column with ten tiers of small cells at the heart sage (45), and is much reduced in size and shape compared with SRB and CB suspensors (Fig. 1*B*, *C*, and *E*) (13). We used LCM to capture SB and *Arabidopsis* embryo proper and suspensor regions (Fig. *1M* and *N*), sequenced each mRNA population using RNA-Seq, and selected for >5-fold up-regulated embryo-proper- and suspensor-specific mRNA sets, including those encoding TFs (Fig. 7*A* and *B*, and Dataset S1). We obtained 95% overlap with *Arabidopsis* embryo proper and suspensor mRNA sequences identified by us previously using GeneChip technology and the same initial cDNAs (Fig. 7*C*) (33) (see Materials and Methods). In addition, there was 82% overlap with suspensor nuclear RNA sequences generated by others using RNA-Seq, indicating that we have a good representation of *Arabidopsis* globular embryo region mRNAs (46).

**Fig. 7.**
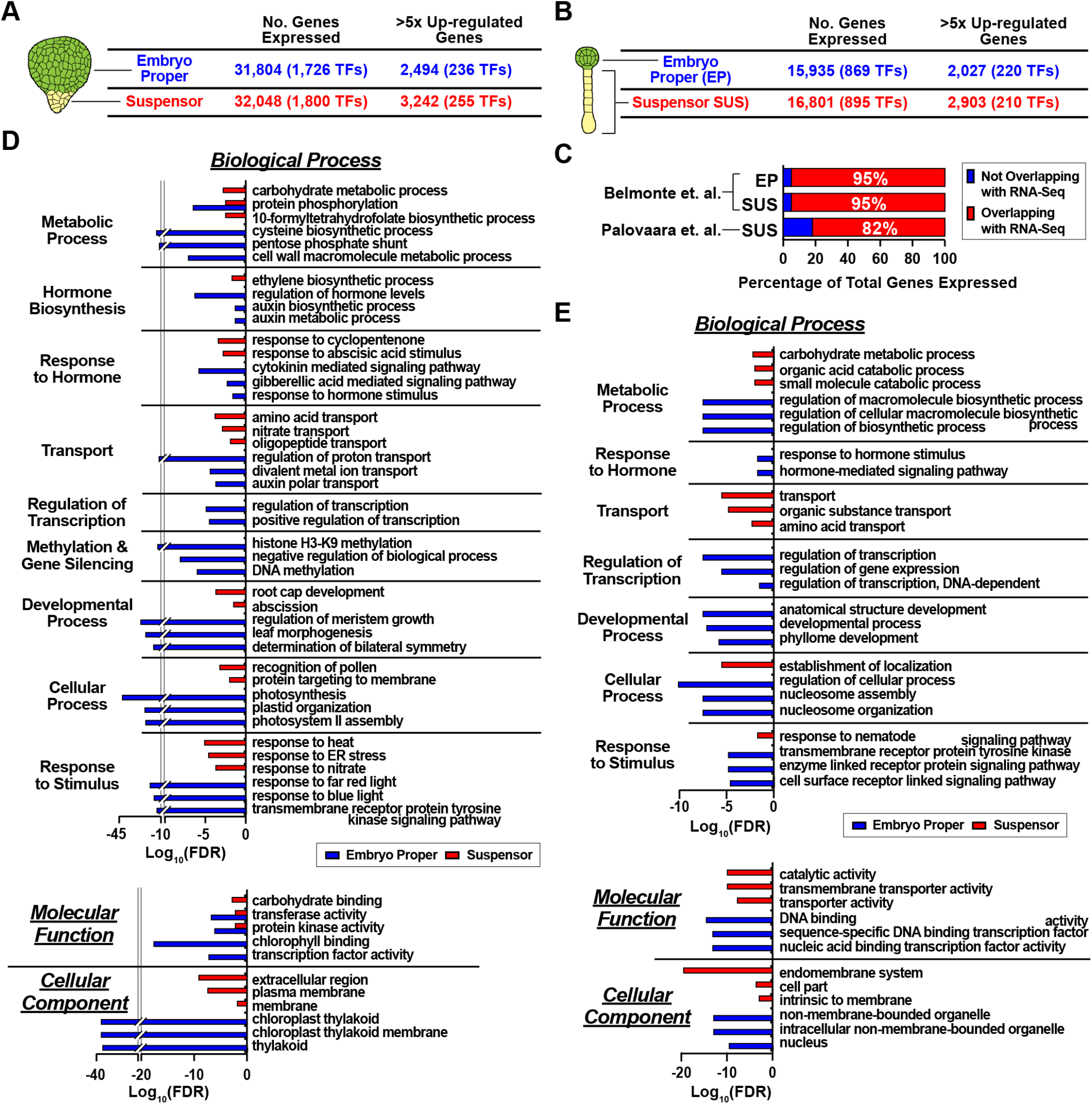
Gene activity in SB and *Arabidopsis* suspensor and embryo proper regions. (*A* and *B*) SB (*A*) and *Arabidopsis* (*B*) genes active (>0.5 RPKM) in at least one biological replicate and 5 fold up-regulated genes in embryo proper and suspensor (see Materials and Methods). (*C*) Percentage of *Arabidopsis* embryo proper and suspensor mRNAs that overlap with datasets published previously by our lab (33) and others (46). (*D* and *E*) Enriched biological process, cellular component, and molecular function GO terms for up-regulated SB (*D*) and *Arabidopsis* (*E*) embryo proper and suspensor mRNAs. Only the top three GO terms for SB and *Arabidopsis* are listed. All expressed genes and those up-regulated >5-fold are listed in Dataset S1. GO terms are listed by functional categories in Dataset S2. TFs, transcription factors; FDR, false discovery rate.

Each mRNA population had a wide range of prevalences, including TF mRNAs and >5-fold region-specific sets (*SI Appendix* Fig. S12), analogous to what was observed with SRB and CB embryo mRNA populations (Fig. 2*B* and *SI Appendix* Fig. S9*B*). Significantly, highly prevalent GA biosynthesis mRNAs were not present in the *Arabidopsis* and SB suspensor up-regulated mRNA sets (*SI Appendix* Fig. S12), in marked contrast with SRB and CB mRNA suspensor populations (Fig. 2*B* and *SI Appendix* Fig. *S9B*). We generated GO terms using the up-regulated SB and *Arabidopsis* embryo proper and suspensor mRNAs (Fig 7*D* and *E*, and Dataset S2). The spectrum of GO terms for the SB and *Arabidopsis* embryo proper regions was similar to those obtained with SRB and CB (Fig. 4 and *SI Appendix* Fig. S10). For example, among the most significant embryo proper GO terms were those involved in regulation of transcription, developmental processes, and response to hormone stimulus, among others (Fig. 7*D* and *E*). By contrast, GO terms obtained with SB and *Arabidopsis* suspensor regions differed significantly from those obtained with SRB and CB (Fig. 4 and *SI Appendix* Fig. S10). The most significant SB and *Arabidopsis* GO terms, such as those for carbohydrate metabolic process and transport (Fig. *7D* and *E*), were similar to those obtained with SRB and CB suspensor mRNAs (Fig. 4 and *SI Appendix* Fig. S10). Missing, however, were GO terms for specialized metabolic processes and pathways, such as oxidation reduction process, glycolysis, pentose phosphate shunt, MEP pathway, and several hormone biosynthesis pathways (e.g., GA and JA), among others, which were hallmarks for SRB and CB suspensors (Fig. 4 and *SI Appendix* Fig. S10). Together these data indicate that, on a functional level, the spectrum of processes carried out by the embryo proper region of all plants investigated was similar irrespective of embryo size and morphology. However, major differences occurred between the small and relatively simple *Arabidopsis* and SB suspensor regions on the one hand, and the giant, specialized SRB and CB suspensors on the other.

### Identification of a Set of Shared Embryo Proper and Suspensor TF mRNAs

We searched the up-regulated embryo proper and suspensor populations for TF mRNAs that were specific to the embryo regions of all plants investigated (Fig. 8 and *SI Appendix* Table S2). For this comparison we used a stringent criterion that included (i) >5-fold up-regulation and (ii) concordance between all biological replicates (see Materials and Methods). We obtained two small sets of globular stage embryo-proper- and suspensor-specific TF mRNAs that were up-regulated in SRB, CB, *Arabidopsis,* and SB embryo regions (Fig. 8*A* and *B*, and *SI Appendix* Table S2). Suspensor TF mRNAs included the known regulators WOX8/9, HDG11, and ARF16 (Fig. 8*A*) (12). On the other hand, embryo proper TF mRNAs included those encoding CUC2, HANABA TANARU (HAN), and TARGET OF MONOPTEROS 5-LIKE 1 (TMO5L1), among others (Fig. 8*B*). These TF mRNAs are involved in shoot meristem, root pole, and vascular tissue differentiation processes, within the embryo proper, respectively (12). STM TF mRNA was specific for SRB, CB, and SB embryo proper regions, but was absent for unknown reasons in the *Arabidopsis* embryo-proper-specific mRNA set, although it was up-regulated within our *Arabidopsis* GeneChip population (33).

**Fig. 8.**
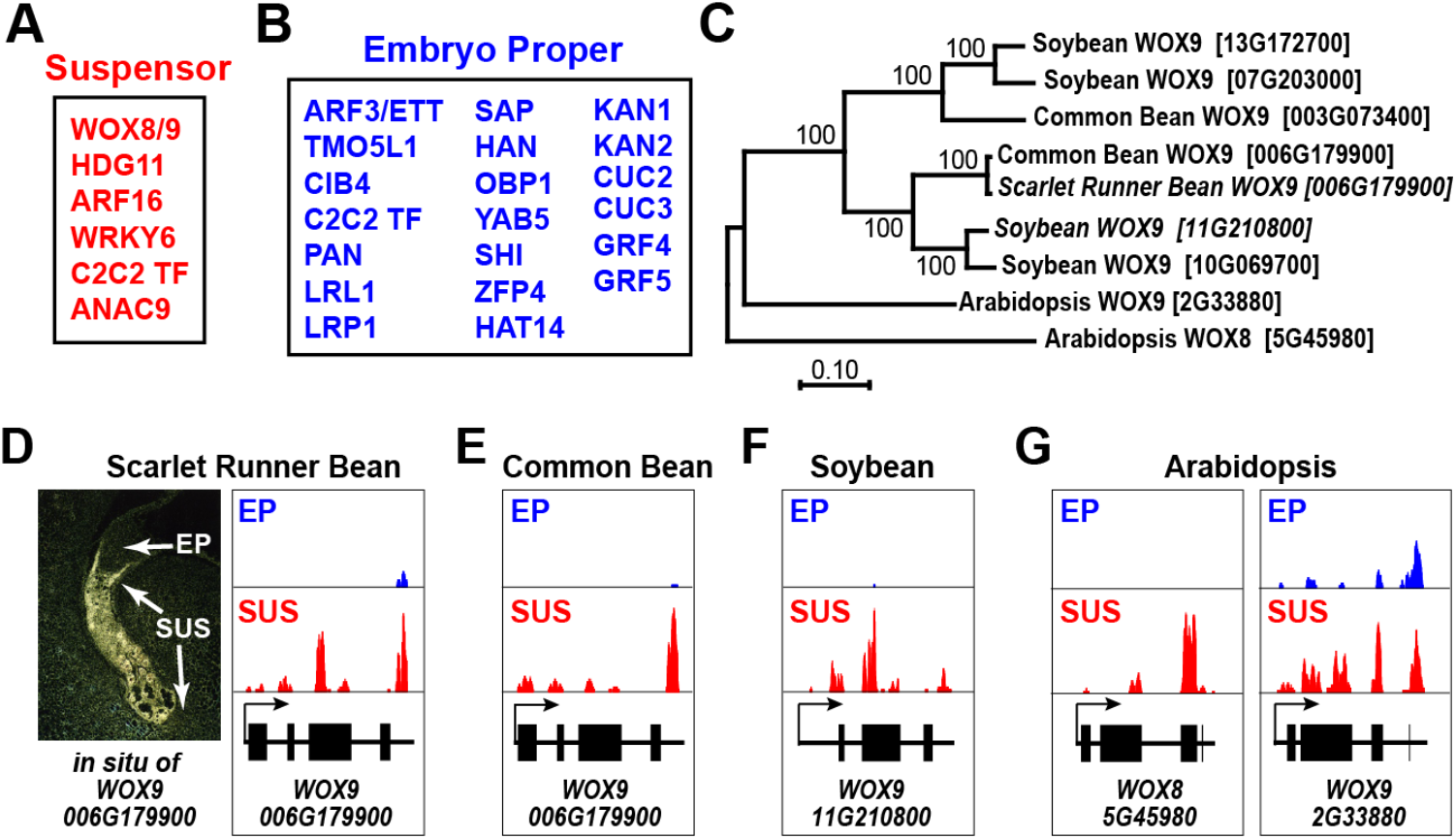
Transcription factor mRNAs that are up-regulated in the suspensor and embryo proper of all four plant species investigated. (*A* and *B*) Shared up-regulated transcription factor mRNAs in the suspensor (*A*) and embryo proper (*B*). Gene abbreviations and IDs are listed in *SI Appendix* Tables S1 and S2, respectively. (*C*) Relationships between SRB, CB, SB, and *Arabidopsis WOX9* genes. The phylogenetic trees were constructed using MEGA X based on the protein sequence alignments with the maximum likelihood method and default parameters (76). Bootstrap confidence values are shown as percentages on each branch. SB (Glyma.11G210800) and SRB (Phvul.006G179900) *WOX9* genes used for ChIP-Seq experiments (Fig. 9) are in italics. (*D-F*) Representation of WOX9 mRNAs in SRB (*D*), CB (*E*), and SB (*F*). SRB WOX9 mRNA *in situ* hybridization image in (*D*) was published previously by our laboratory as EST PCEP0357 using author reuse permissions given by the American Society of Plant Biology (29). (*G*) Representation of *Arabidopsis* WOX8 and WOX9 mRNAs.

### Several WOX9 TF Targets are Shared By SRB and SB

*Arabidopsis WOX8,* and its close relative *WOX9,* play important roles in suspensor differentiation (12). *WOX8/9* gene relatives in SRB, CB, and SB most closely resembled *Arabidopsis WOX9* by phylogenetic analysis (Fig. 8*C*), but *WOX8* by their suspensor-specific expression patterns (Fig. 8 *D-G*) (12). We named the SRB, CB, and SB relatives as *WOX9,* and considered these genes as functionally equivalent to *Arabidopsis WOX8*.

We carried out ChIP-Seq experiments with SRB and SB WOX9 peptide antibodies to (i) uncover WOX9 downstream target genes, (ii) characterize target functions, and (iii) ascertain whether any targets were shared between SRB and SB (Fig. 9) (see Materials and Methods). We used whole globular-stage seeds for our experiments, because in both plants (i) WOX9 was the most prevalent suspensor-specific TF mRNA (Fig. 2*B* and Fig. 3*B*, *SI Appendix* Fig. S12), and (ii) only present within suspensor region of the seed [Fig. 8*D* and Harada-Goldberg LCM datasets (seedgenenetwork.net)]. We obtained 660 and 178 potential WOX9 target genes in SB and SRB, respectively, including 88 and 21 TF gene targets (Fig. 9*A* and Dataset S4). We defined target genes as those that were at the intersection of WOX9 bound genes and genes up-regulated >5-fold within the suspensor (see Materials and Methods) (47). We carried out GO term analysis on the SB and SRB WOX9 target genes (Fig. 9*B*). The most significant GO term in both plants was regulation of transcription, reflecting the large representation of TF genes in SB and SRB target gene sets (Fig. 9*A*). Significantly, the range of GO terms for SRB WOX9 targets was greater than those for SB, and included GO terms that mirrored many functional activities unique to the SRB suspensor region (Fig. 4). These included oxidation reduction processes and JA biosynthesis, among others (Fig. 9*B* and Dataset S4), but did not include GA biosynthesis.

**Fig. 9.**
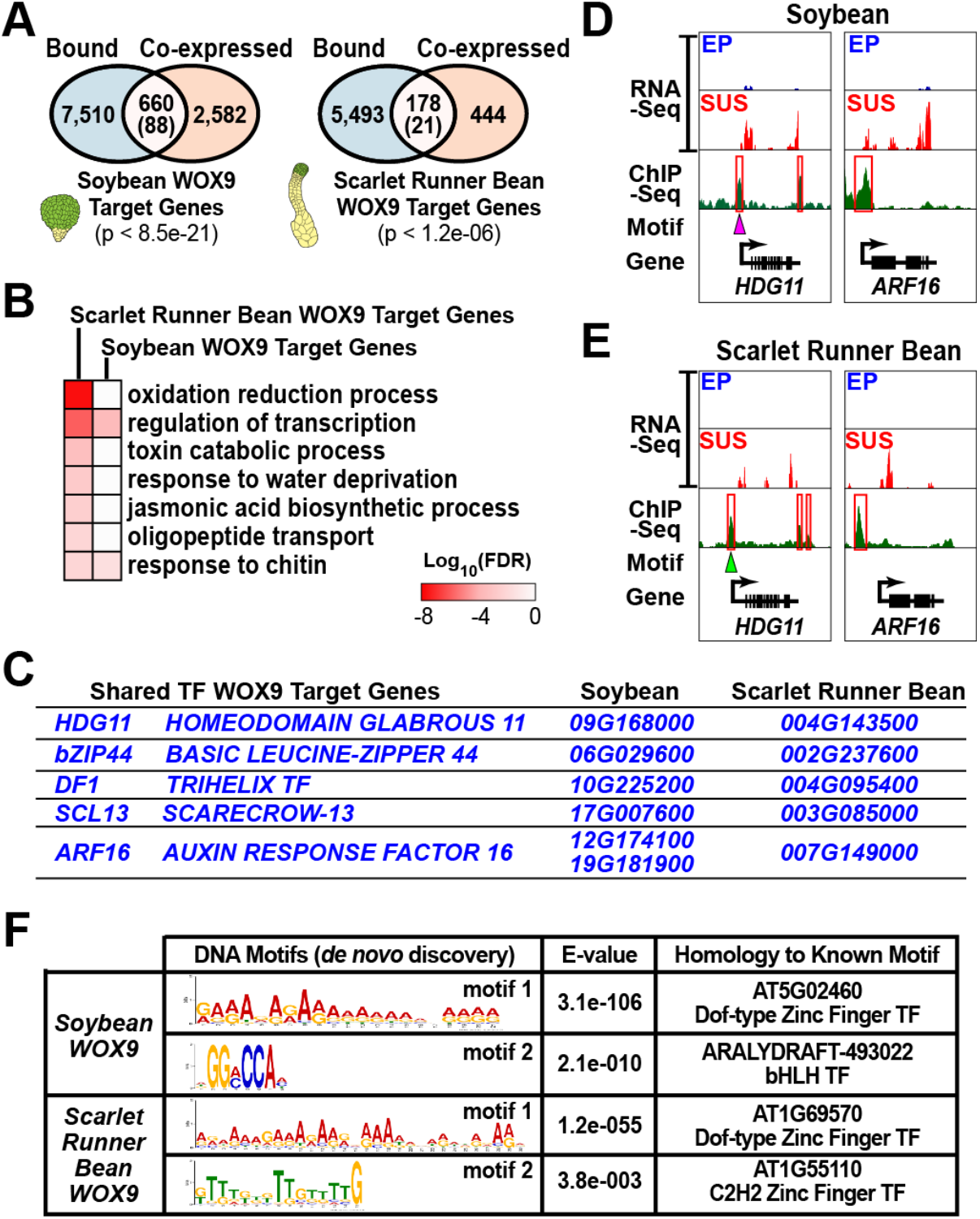
WOX9 transcription factor targets in SRB and SB globular stage seeds. (*A*) Venn diagrams between WOX9 bound genes (blue) and co-expressed genes (orange). Bound genes are those with peaks within 1 kb upstream of the transcription start site (47). Co-expressed genes are the >5-fold up-regulated genes shown in Fig. 2 (SRB) and Fig. 7 (SB). WOX9 target genes are assumed to be those in the intersection between bound and co-expressed genes (pink) (47, 65). TF genes are in parentheses. P values were obtained using a hypergeometric distribution test (65). WOX9 bound regions and target genes are listed in Dataset S4. (*B*) The most significant GO terms for SRB and SB WOX9 target genes. All GO terms are listed in Dataset S4. (*C*) WOX9 TF target genes that are shared by SRB and SB (see Materials and Methods). (*D* and *E)* Genome browser view of shared SB (*D*) and SRB (*E*) WOX9 target gene activity (RNA-Seq) and bound gene regions (ChIP-Seq). Arrows point to motif 1 in both SB and SRB *HDG11* genes. *ARF16* did not have an enriched motif. (*F*) Enriched motifs in target genes bound by WOX9. The MEME-ChIP suite (66) was used for *de novo* motif discovery as previously described (47, 65). The E-value is the probability of obtaining a specific motif compared with a randomly generated set of sequences (66). The Tomtom tool and plant TF databases within the MEME-ChIP suite were used to identify TFs with binding sites similar to the discovered motifs (77–79). The zinc finger TFs were present in the *Arabidopsis* DAP-Seq database (78), whereas the bHLH TF was identified from the *Arabidopsis* protein-binding microarray database (79). Target genes associated with each enriched motif are listed in Dataset S4.

We searched for TF target genes that were shared by SB and SRB, and found a small number that included *HDG11* and *ARF16* genes that were present in the suspensor-specific gene set that was shared by all plants investigated (Fig. 8*A* and Fig. 9*C*). RNA-Seq and ChIP-Seq genome browser views show (i) the suspensor-specific expression pattern of SB (Fig. 9*D*) and SRB (Fig. 9*E*) *HDG11* and *ARF16* genes, and (ii) indicate that WOX9 binding peaks were near their transcription start sites (Fig. 9*D* and *E*). We carried out motif analysis on the SB and SRB WOX9 targets and uncovered two distinct binding motifs for SB and SRB (Fig. 9*F*). Approximately 30% of SB and SRB target genes had one, or both, of these motifs in their upstream regions. One SB motif recognized a Dof-type zinc finger TF (motif 1), while the other a beta helix-loop-helix TF (motif 2) (Fig. 9*F*). By contrast, both SRB motifs recognized Dof-type zinc finger TFs, and were similar to the SB Dof-type zinc finger motif (Fig. 9*F*). One candidate binding to this motif might be the Dof-type zinc finger TF that was up-regulated in all suspensor mRNA populations we investigated (Fig. 8, *SI Appendix* Table S2, and Dataset S1). Together, these data suggest that (i) WOX9 binds to a large number of potential target genes with distinct overall functions in SB and SRB suspensors, (ii) several TF gene targets are shared between SB and SRB, and (iii) WOX9 appears to form a complex with other TFs in order to bind to a subset of target genes.

## Discussion

We compared the embryo proper and suspensor transcriptomes of SRB, CB, SB, and *Arabidopsis* globular-stage embryos. At this stage of development, major regulatory decisions are being made, particularly within the embryo proper region (12), and embryos of these species differ in size and morphology, reflecting their final seed sizes and differences in suspensor morphology (Fig. 1). Our goals were to (i) describe the major developmental and metabolic processes that occur within embryo proper and suspensor regions at the globular stage, (ii) characterize the functional differences between giant specialized suspensors and structurally simpler suspensor regions, (iii) uncover sets of TFs that are shared between embryo proper and suspensor regions, irrespective of morphology, which might play essential roles in region-specific differentiation events, and (iv) begin to dissect suspensor genetic regulatory networks by investigating the downstream targets of an essential suspensor regulator – WOX9 – in both SRB and SB.

### Giant SRB and CB Suspensors Are Specialized to Express Specific Metabolic Pathways

A comparison of major GO terms derived from up-regulated suspensor mRNAs shows the dramatic functional differences between SRB and CB suspensors with those of SB and *Arabidopsis* (Fig. 4, Fig. 7, *SI Appendix* Fig. S10, and Dataset S2), and are summarized conceptually in Fig. 10. Significantly, a large number of metabolic pathways leading to the synthesis of several important hormones are overrepresented in SRB and CB suspensor-specific mRNAs. Up-regulated mRNAs are also enriched in plastid organelle processes (Fig. 10), as well as other plastid-associated GO terms such as starch biosynthesis, plastid stroma, and plastid membrane, because many of the major pathways are localized within specialized plastids (Dataset S3). Remarkably, SRB and CB suspensor-specific mRNAs are overrepresented in a continuum of biosynthetic pathways starting with glycolysis, pentose phosphate shunt, and the MEP isoprenoid pathway on one hand, and ending with major metabolic pathways leading to the production of GA, JA, and auxin, among others (Fig. 4, *Appendix SI* Fig. S5 and Fig. S10, and Dataset S2). The machinery driving these pathways is active, because their end-product hormones are present in SRB suspensors (24, 27, 26, 39). SRB and CB up-regulated mRNAs are also overrepresented in the transport and signaling processes making it possible for hormones (e.g., GA, JA, and auxin) synthesized in the suspensor to be either utilized within the suspensor region or exported to the embryo proper to facilitate developmental and physiological events (Fig. 4, Fig. 10, *SI Appendix* Fig. S10, and Dataset S2).

**Fig. 10.**
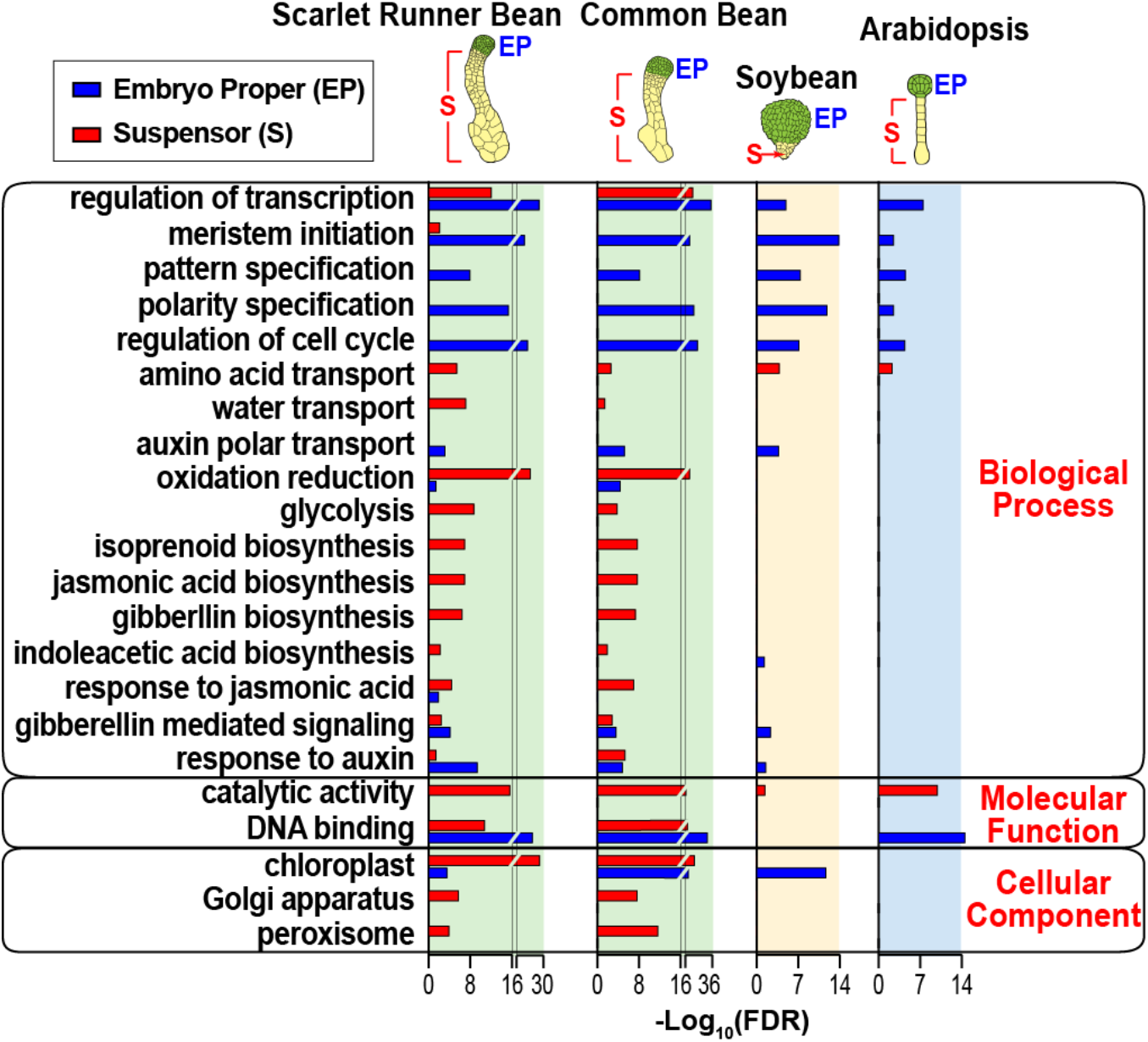
Comparison of SRB, CB, SB, and *Arabidopsis* GO terms from up-regulated suspensor and embryo proper mRNAs. Summary of the most significant shared and specific GO terms. Data taken from Dataset S2, Fig. 4, Fig. 7, and *SI Appendix* Fig. S10.

With the exception of auxin, suspensor-specific mRNAs are not overrepresented in GO terms for the majority of pathways leading to hormone production (e.g., MEP pathway, glycolysis, pentose phosphate shunt) in simpler SB and *Arabidopsis* suspensors (Fig. 7, Fig. 10, and Dataset S2). Nor are up-regulated mRNAs enriched for plastid-associated GO terms in these plants (Fig. 10). Where then are hormone synthesizing processes being carried out in SB and *Arabidopsis* seeds? We previously showed that *Arabidopsis* chalazal-endosperm-specific mRNAs are overrepresented for GA, ABA, CK, and auxin GO terms at the globular stage – suggesting that these hormones are produced within this specialized endosperm subregion and not in the suspensor (33). We searched the Harada-Goldberg SB seed LCM datasets and found that these hormones are most likely produced in globular stage endosperm and seed coat regions (seedgenenetwork.net) (47). For example, the only the seed region that has all of the mRNAs required for GA biosynthesis, including GA3ox, is the outer integument seed coat layer, and none of the rate limiting enzymes for JA, ABA, CK, and auxin are up-regulated within the suspensor. Thus, highly specialized SRB and CB suspensors may have co-opted regulatory events that occur within the endosperm and seed coat layers of plants that have simpler suspensors, such as SB and *Arabidopsis* – a hypothesis that was proposed over 100 years ago (5, 9, 27). Whether pathways leading to similar hormones in distinct seed parts utilize the same or different gene regulatory pathways remains to be determined.

The high degree of metabolic functional specialization within giant SRB and CB suspensors reported here is in remarkable agreement with elegant histological studies carried out by Ed Yeung with SRB suspensors over four decades ago (22). His experiments revealed that the SRB suspensor has a large number of specialized starch-containing plastids, extensive networks of wall ingrowths, and substantial amounts of smooth membranes and dictyosomes consistent with (i) an embryo region that is synthesizing and transporting essential materials to the embryo proper, and (ii) the GO terms we uncovered that are generated by up-regulated suspensor mRNAs. Finally, are the giant suspensor regions of other plants specialized for hormone production as well? *Cytisus laburnum,* or Golden Chain, and *Tropaeolum majus,* the Garden Nasturtium, both have giant suspensors and have been shown to synthesize bioactive GAs (48). Thus, the development of giant specialized suspensors appears to be associated with specialized metabolic processes such as those leading to hormone production. How plants such as SRB and CB evolved morphologically unique suspensor regions as well the specialized regulatory processes that drive and coordinate the expression of large numbers of specific metabolic pathway genes remains an important unanswered question.

### The Embryo Proper Region Carries Out Similar Processes in Plants With Different Suspensor Morphologies

In contrast with the suspensor, the constellation of functions carried out by the embryo proper regions of SRB, CB, SB, and *Arabidopsis* are similar – irrespective of suspensor morphology (Fig. 4, Fig. 7, SI Appendix Fig. S10, and Dataset S2). Up-regulated embryo proper mRNAs are enriched for GO terms reflecting gene regulation, epigenetic events, pattern formation, hormone responses, cell proliferation, and DNA replication, among many others (Fig. 10). These GO terms mirror the cell division and differentiation processes that occur within SRB, CB, SB, and *Arabidopsis* globular stage embryo proper regions as they form cells, tissues, and subregions that will constitute the embryo when it matures. Thus, the evolutionary events that give rise to morphologically diverse suspensor regions that degenerate during seed development are uncoupled from those that maintain continuity of embryo proper form and function in order to guarantee plant survival from generation to generation.

### TF mRNA Sets Have Been Identified That Are Shared By Embryo Proper and Suspensor Regions Irrespective of Embryo Morphology

#### Suspensor-specific TF mRNAs

We uncovered a small number of mRNAs that are up-regulated in SRB, CB, SB, and *Arabidopsis* suspensor regions (Fig. 8*A*). The precise role that most of these TF mRNAs play in suspensor differentiation and function is not yet known. However, most likely they function within all suspensor cells as localization experiments carried out with suspensor-specific mRNAs [e.g., WOX9 (Fig. 8*D*) and GA enzymes], by us (28, 30), and by others (11, 49), showed that transcripts are distributed relatively evenly across the suspensor. The absence of cellular diversity suggests that genetic regulatory networks responsible for controlling suspensor form and function are probably simpler than those that operate in the embryo proper which undergoes a more complex set of developmental events required to establish the diverse cell types and tissues of this embryonic region.

WOX8/9, HDG11, and WRKY2 TF mRNAs have been shown to play essential roles in *Arabidopsis* suspensor differentiation (11, 50). WRKY2 mRNA is up-regulated >5-fold in our *Arabidopsis* suspensor mRNA population (Dataset S1). Close relatives in SRB (Phvul.005G005800; Phvul.008G054100), CB (Phvul.005G005800; Phvul.008G054100), and SB (Glma.09250500) suspensor mRNAs are up-regulated 3 to 6-fold depending upon the gene (Dataset S1), but collectively failed to meet our >5-fold criterion for shared mRNAs, in contrast with WOX8/9 and HDG11 TF mRNAs (Fig. 3*C*, Fig. 8, and *SI Appendix* Fig. S9*G*). Both WRKY2 and HDG11 TFs are essential for *WOX8/9* gene activation within the suspensor, and are part of the SHORT SUSPENSOR (SSP)/YODA signaling cascade required for suspensor differentiation (11, 50). HDG11 mRNA is present in the egg cell, suggesting that maternal factors play a role in suspensor specification and are important for *WOX8/9* gene activation (11). We proposed over two decades ago that localized maternal factors in the egg cell might be distributed asymmetrically to the basal cell after zygote division promoting suspensor differentiation similar to maternal localization processes that occur in animal embryos such as the sea urchin (30). HDG11 TF mRNA localization within the egg cell and its role in suspensor development is consistent with this hypothesis. Nevertheless, our results suggest that WRKY2, HDG11, and WOX8/9 TF mRNAs perform similar roles in SRB, CB, and SB, and that regulatory events giving rise to the suspensor early in embryogenesis are conserved in plants – irrespective of suspensor morphology.

#### Embryo-proper-specific TF mRNAs

We uncovered a larger number of globular stage embryoproper-specific TF mRNAs that are shared between SRB, CB, SB, and *Arabidopsis* (Fig. 8B), most likely a result of the greater complexity of the embryo proper region. Many of these TF mRNAs have known roles in early embryo proper development as a result of the elegant genetic experiments carried out with *Arabidopsis* (12, 51). What emerges from these studies is that the globular embryo is divided into specific territories marking specification events leading to shoot and root meristems, vasculature regions, cotyledons, and other subregions and tissues of the mature embryo. Many of these territories are generated by auxin gradient signaling that sets off a cascade of events leading to the differentiation of specific embryo parts and subregions (12). In this respect, the embryo proper region resembles conceptually an early sea urchin embryo that is divided into specific territories that require different regulatory inputs to specify unique differentiation events leading to the mature embryo (52). The up-regulation of shared TF mRNAs in the embryo proper of SRB, CB, SB, and *Arabidopsis* suggests that common regulatory pathways operate within the embryo proper across the plant kingdom to define analogous specification domains. In contrast with the suspensor, it will be a significant challenge to unravel the mosaic of genetic regulatory networks that govern differentiation events within each embryo proper territory. The precise architecture of these networks and how they program the differentiation of a mature plant embryo remain to be determined.

### Gaining Entry Into WOX9 Suspensor Regulatory Networks

One of the interesting aspects of the WOX9 ChIP-Seq experiments in both SRB and SB is that there is a significant enrichment in TF target genes. This is consistent with the essential role that *WOX9* plays in suspensor differentiation (11), and suggests that *WOX9* is upstream in the hierarchy of suspensor regulatory networks. It is surprising that the *HDG11* TF gene appears to be a WOX9 target. This suggests that there is a feedback loop in the regulatory circuit containing *WOX9*, and that after HDG11 activation of *WOX9,* WOX9 plays a role in reinforcing *HDG11* transcription. What is interesting is that both HDG11 and WOX8/9 mRNAs are localized within the *Arabidopsis* egg cell (11), and that the maternally-derived *HDG11* allele is expressed early in suspensor development (53). This suggests that the *WOX9* regulatory network might be activated during egg cell development and sets off a cascade of events leading to suspensor differentiation following fertilization.

What is also significant is the divergence of WOX9 target gene functions in SRB and SB (Fig. 9). These functional differences correlate with morphological differences between SRB and SB suspensor regions (Fig. 1). For example, GA biosynthesis is a key marker of giant specialized suspensors (Fig. 10), but GA genes are not direct targets of WOX9 (Fig. 9 and Dataset S4). How are these genes activated in SRB and CB suspensors? Previously, we uncovered a *cis* regulatory module (CRM) that is required for *GA20ox* and *G564* transcription within the suspensor (28, 31). Both of these genes are up-regulated in SRB and CB suspensor regions (Fig. 2 and *SI Appendix* Fig. S9). This module contains three distinct *cis*-elements, designated as the 10-bp motif, Region 2 motif, and the Fifth motif, which have sequences that recognize Dof-type zinc finger, myb, and C2H2 zinc finger TFs, respectively (28). We searched SRB WOX9 targets for mRNAs encoding these TFs and found candidates for all three that are up-regulated in the SRB suspensor (Datasets S1 and S4). Thus, one model is that activation of WOX9 leads to the activation of TFs which switch on GA biosynthesis and other specialized suspensor-specific genes that are downstream of WOX9 in the regulatory circuitry (*SI Appendix* Fig. S13).

It would appear, therefore, that WOX9 plays a dual role – one that is responsible for basic developmental events required for suspensor differentiation across the plant kingdom, and another that has been adapted for highly specialized functions in giant suspensors such as those in SRB and CB. The regulatory circuitry controlling these specialized suspensor functions has evolved over the 19 mya since the separation of SRB and SB from their common ancestor (Fig. 1*J*). Clearly, the major tasks before us are to (i) unravel the basic regulatory circuitry required for the differentiation of all suspensors irrespective of morphology, and (ii) determine how regulatory genes in that network (e.g., WOX9) are utilized differently in circuits that control specialized downstream suspensor functions in plants with morphologically distinct suspensor regions. What these circuits are and how they are organized within plant genomes remain to be discovered.

## Materials and Methods

### Plant Growth

#### SRB

Detailed methods for growing and using SRB for embryo genomics experiments were published recently by our lab (20). In brief, SRB *(Phaseolus coccineus* cv Hammond’s Dwarf Red Flower) (Vermont Bean Seed Company) (54) was grown in a greenhouse with a 16 h/8 h day/night cycle at 22° C to 30° C. Plants flowered approximately one month after seed germination and open flowers were hand-pollinated (20). Pods were harvested 5-7 days after pollination (DAP) to obtain globular stage embryos from seeds 2.0 to 2.5 mm in length (Fig. 1*A-C*).

#### CB

*Phaseolus vulgaris* (Andean inbred landrace accession G19833) seeds were planted and CB plants were grown under the same conditions as SRB (20), except after one month they were moved into a growth chamber with an 8 h/16 h light/dark cycle for one week to induce flowering. Pods from self-pollinated flowers were collected at 5-6 DAP to harvest seeds 1.6 to 2.0 mm in length containing globular stage embryos (Fig. *1D* and *E*).

#### SB

*Glycine max* (cv Williams 82) was grown in the greenhouse, and seeds 1.0 to 1.5 mm in length containing globular stage embryos (Fig. 1*F* and *G*) were harvested as described in detail previously (47, 55).

#### Arabidopsis

*Arabidopsis thaliana* [ecotype Wassilewskija (Ws-0)] plants were grown in the greenhouse as described previously (56). Seeds from siliques 1.0-1.1 cm in length containing globular stage embryos (Fig. 1*H* and *I*) were collected three to four DAP (33).

#### SRB Genome Sequencing and Assembly

Genomic DNA was isolated from SRB leaves using the DNeasy Plant Mini Kit (Qiagen) according to the manufacturer’s protocol. A sequencing library was constructed using the Illumina TruSeq DNA Sample Prep Kit. Single-end and paired-end 50 bp reads were generated using an Illumina HiSeq 2000. The SRB genome was assembled using the short read *de novo* assembler ABySS (57). Contigs >200 bp in length were deposited in GenBank as contigs QBDZ01000001 to QBDZ01192921.

#### LCM

Procedures used in our laboratories for fixing, embedding, and capturing seed sections by LCM have been described in detail previously (33). The same methods were used for all four plant species discussed in this paper. In brief, seeds containing mid-stage globular embryos (Fig. 1) were fixed, embedded in paraffin, and sliced into 6 μm sections. Embryo proper and suspensor regions were captured using a Leica LMD 6000 Microdissection System (Leica Microsystems) (Fig. *1K-N*). Only medial sections containing entire globular stage embryos were used for LCM to avoid contamination from surrounding endosperm and seed coat tissues (20) (Fig. 1*K*). Two to three biological replicates were captured for each embryonic region, except for *Arabidopsis* as there was only enough material left from one of the two biological replicates used previously in our GeneChip seed gene expression experiments (33). Captured materials were collected on the cap of a 0.2 mL microcentrifuge tube containing 30 μl of PicoPure RNA Isolation Kit lysis solution (Thermo Fisher Scientific) and stored at −70° C prior to total RNA isolation.

#### RNA Isolation

Total RNA from laser captured embryo proper and suspensor regions was isolated using the PicoPure RNA Isolation Kit (Thermo Scientific) according to the manufacturer’s instructions.

#### RNA-Seq Library Construction and Sequencing

SRB, CB, and SB sequencing libraries were prepared using 5 ng of total RNA from suspensor and embryo proper laser captured regions. Double-stranded cDNA was synthesized and amplified using the Ovation RNA-Seq System V1 (NuGen) according to the manufacturer’s directions. *Arabidopsis* embryo proper and suspensor sequencing libraries were constructed from double-stranded cDNAs that were used previously to generate the GeneChip data presented in our *Arabidopsis* seed gene atlas paper (33). RNA-Seq libraries were prepared using the TruSeq DNA Sample prep kit (Illumina). Single-end 50 bp reads were generated using an Illumina HiSeq 2000 sequencing system.

#### RNA-Seq Data Analysis

RNA-Seq reads were downloaded from the UCLA Broad Stem Cell Center Sequencing Core and subjected to quality filtering. Adapter sequences were removed using the FASTX-toolkit (http://hannonlab.cshl.edu/fastx_toolkit/). DNA sequences passing the Illumina purity filter were selected and subjected to base-call error analysis. Either HISAT2 (version 2.0.6) (58) or Bowtie (version 0.12.7) (59) was used to map sequences to each reference genome: (i) CB *(Phaseolus vulgaris* accession G19833 v. 2.1) (https://phytozome.jgi.doe.gov/) (32); (ii) SB *(Glycine max* cv Williams 82 v. 1.1] (https://phytozome.jgi.doe.gov/); and (iii) *Arabidopsis* (*Arabidopsis thaliana* ecotype Ws-0 v. TAIR10) (https://www.arabidopsis.org). Only reads that mapped to one position within the genome were used for subsequent analysis. Read counts for each gene model were computed using HTSeq (v. 0.6.1) (60) and converted to RPKM values (35). A cutoff of <0.5 RPKM was chosen as the lower limit for transcript representation in each mRNA population (Fig. 2*B*) (36). RNA-Seq data were deposited in the Gene Expression Omnibus (GEO) database with accession numbers: (i) GSE57537 for CB and SRB; (ii) GSE57349 for SB; and (iii) GSE135393 for *Arabidopsis.*

#### Identification of Up-Regulated mRNAs

RNA-Seq reads were normalized by the TMM (Trimmed Mean of M-values) method using EdgeR package v. 3.4.2. (61). Suspensor and embryo proper mRNA populations were compared, and mRNAs that had RPKM values >5-fold higher in each embryo region compared with the other were designated as up-regulated mRNAs (FDR <0.05) (47).

### GO Term Enrichment Analysis

#### *SRB* and *CB*

GO annotations of CB genes are not yet comprehensive enough to provide a meaningful profile of SRB and CB suspensor and embryo proper biological activities. To overcome this problem, we utilized the more extensive SB GO annotations because of the close evolutionary distance (19 mya) between SB, CB, and SRB (Fig. 1*J*). SB homologs of CB genes were identified using the top hits obtained from BLASTP analysis. Overrepresented GO terms were identified in SRB and CB suspensor and embryo proper mRNAs by using Soybase GO annotations (http://soybase.org/genomeannotation/index.php). GO term enrichment analysis was performed using the GoSeq R Bioconductor package (http://www.r-project.org/) (62) using the following parameters: (i) reference library – SB homologs of CB genes; (ii) statistical test method – hypergeometric; and (iii) significance level – Benjamini-Hochberg multiple testing correction (FDR <0.05).

#### SB

SB suspensor and embryo proper GO term analysis was carried out using Soybase GO annotations (http://soybase.org/genomeannotation/index.php) following the methods used for SRB and CB.

#### Arabidopsis

*Arabidopsis* suspensor and embryo proper GO enrichment analysis was carried out using the VirtualPlant 1.3 program (http://virtualplant.bio.nyu.edu/cgi-bin/vpweb/) (63) using FDR cutoff value <0.05 (Benjamini–Hochberg multiple testing correction).

#### Identification of Enzyme Subcellular Locations

The Plant Metabolic Pathway Database (v. 13) (https://www.plantcyc.org) was used to identify metabolic pathways encoded by up-regulated suspensor and embryo proper enzyme mRNAs. Enzyme protein sequences were used to predict their subcellular locations using the machine learning program DeepLoc-1.0 (64).

#### Identification of Conserved Up-Regulated TF Genes

*Arabidopsis* homologs of ≥ five-fold up-regulated SRB, CB, and SB suspensor and embryo proper genes were obtained based on their annotations generated by Phytozome (https://phytozome.jgi.doe.gov). The top hit for each gene was considered to be the most closely related *Arabidopsis* homolog.

#### ChIP-Seq Experiments

SRB and SB seeds containing globular stage embryos were used to identify WOX9 TF gene targets (Fig. 1 *B* and *G*). ChIP assays, library construction, DNA sequencing analysis, and peak calling were carried out using ENCODE guidelines as described elsewhere by our laboratories (47, 55, 65) – including: (i) cross correlation analysis for data quality; (ii) irreproducible discovery rate (IDR) for replicate consistency; and (iii) MACS2 for peak calling using an IDR threshold of 0.01. WOX9 peptide antibodies used for ChIP assays were synthesized by Eurogentec (https://www.eurogentec.com/en/) using amino acid sequences specific for SRB homolog Phvul.006G179900 and SB homolog Glyma.11G210800 (Fig. 8*C*). The sequences of these peptides were: (i) SRB – PNSPTTSVNQTYFQ and (ii) SB – NHHYPTSLPQTATTA. Two biological replicates were carried out for each ChIP assay, and sequencing data were deposited in GEO as GSE153644 (SRB) and GSE135267 (SB). The MEME-ChIP suite (66) was used for *de novo* motif discovery as previously described (47, 65).

## Author contributions

R.B.G., M.C., B.H.L., A.Q.B., M.P., and J.J.H. designed research; X.W., N.R.A., K.F.H., B.H.L, A.Q.B., J.P., and M.C. performed wet bench research; M.C., J.-Y.L., and S.C. analyzed sequencing data; and R.B.G. and M.C. wrote the paper

## Data Deposition

The transcriptome and ChIP-Seq data reported in this paper have been deposited in the Gene Expression Omnibus (GEO) database, www.ncbi.nih.gov/geo [Scarlet Runner Bean and Common Bean RNA-Seq (accession no. GSE57537); Soybean RNA-Seq (accession no. GSE57349); *Arabidopsis* RNA-Seq (accession no. GSE135393); Scarlet Runner Bean ChIP-Seq (accession no. GSE153644); and Soybean ChIP-Seq (accession no. GSE135267)]. The genome sequences were deposited into the GenBank database, www.ncbi.nlm.nih.gov/genbank/ [Scarlet Runner Bean (contigs QBDZ01000001 to QBDZ01192921)].

## ACKNOWLEDGMENTS

This research was supported by a grant from the National Science Foundation Plant Genome Program to R.B.G., M.P., and J.J.H.

---

**Fig. S1.**
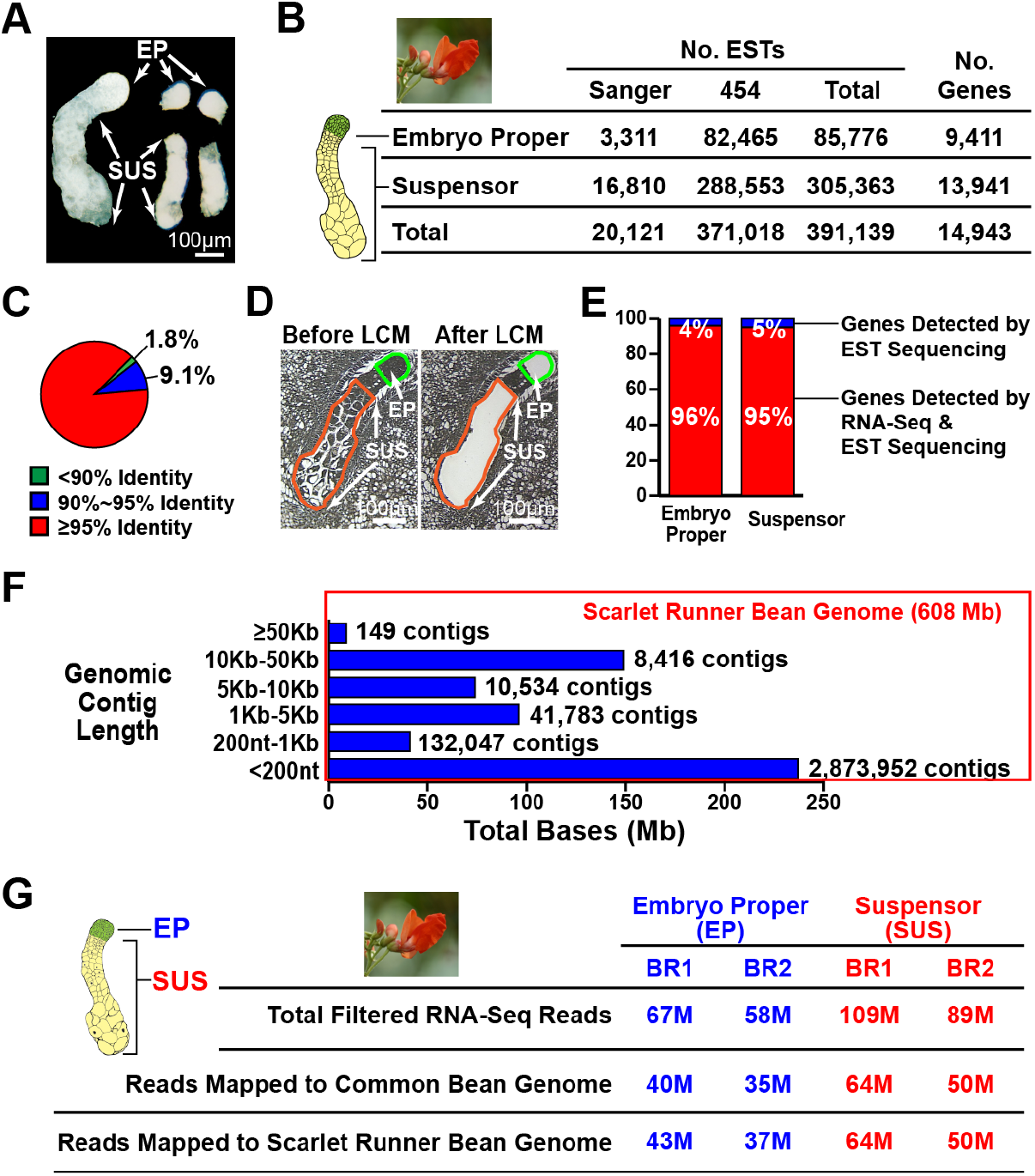
Scarlet Runner Bean (SRB) expressed sequence tag (EST) and genome sequencing. (*A*) Hand-dissected SRB globular stage embryo proper (EP) and suspensor (SUS) regions used for EST sequencing. Procedures for isolating and dissecting SRB embryos were described previously by our laboratory (1). Image was taken from a paper published elsewhere by our laboratory under author reuse permissions given by the American Society of Plant Biology (2). (*B*) ESTs generated from hand-dissected SRB globular stage embryo proper and suspensor mRNAs. Sanger and 454 refer to EST sequencing methods. A small number of Sanger ESTs were described earlier (2). The number of genes represented in EST populations was determined by BLASTN comparisons with predicted gene transcripts from the Common Bean (CB) reference genome (https://phytozome.jgi.doe.gov) (3) (see Materials and Methods). All EST sequences were deposited into GenBank as accession series CA896559 to CA916678 and GD289845 to GD660862. (*C*) Similarities between SRB EST sequences and CB genome predicted transcripts. Circle shows the identity distribution of SRB ESTs and CB genome predicted transcripts taken from BLASTN alignments (E-value <1E-6). (*D*) Laser Capture Microdissection (LCM) of SRB embryo proper (EP) and suspensor (SUS) regions (see Materials and Methods). Image taken from Fig. *1K*. (*E*) Representation of SRB embryo proper and suspensor ESTs (*B*) in RNA-Seq datasets (Fig. 2*A*). *(F)* Distribution of SRB contig lengths obtained from genome sequence assembly (see Materials and Methods). (*G*) Comparison of SRB embryo proper and suspensor RNA-Seq read alignments with SRB and CB genome sequences.

**Fig. S2.**
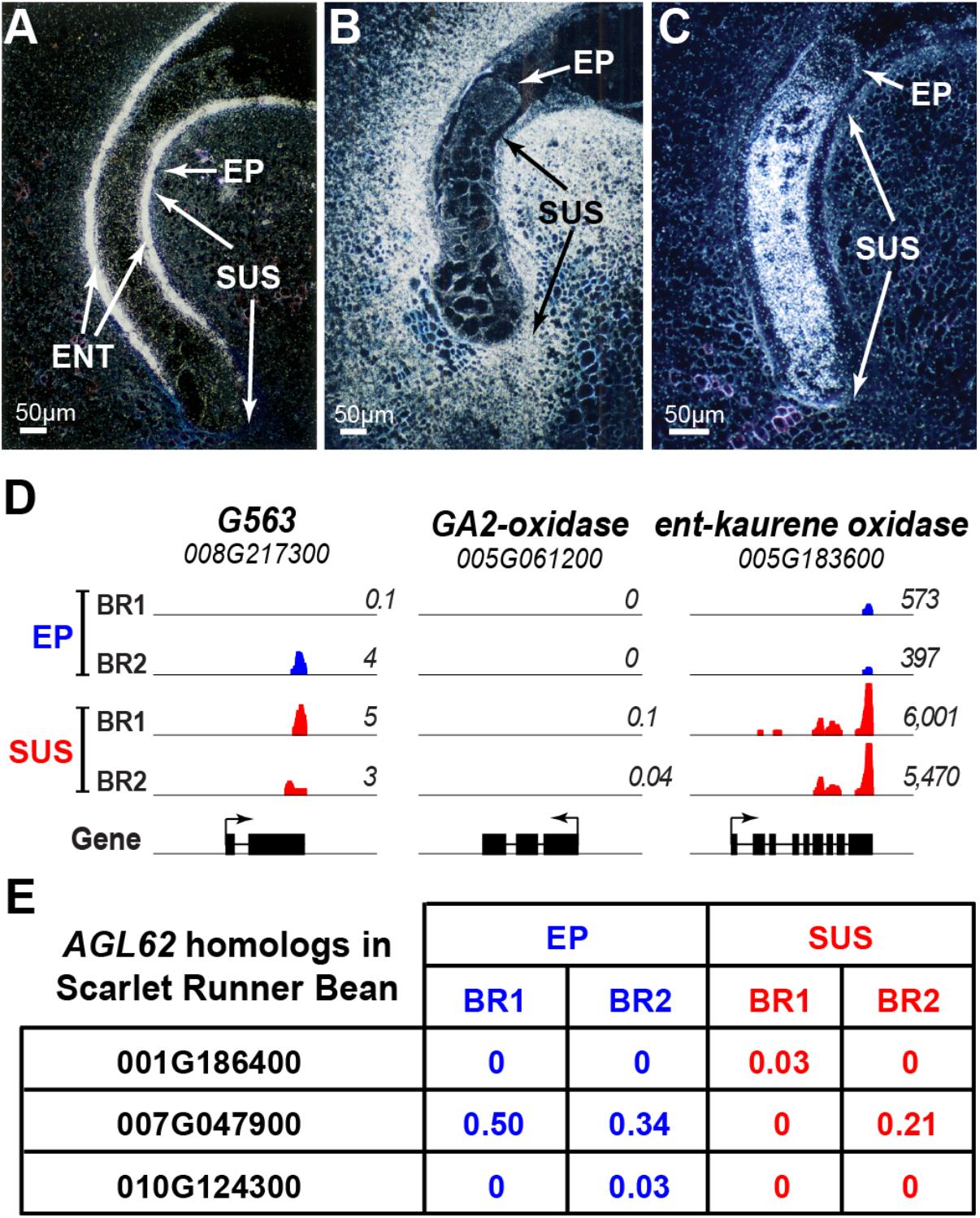
Absence of contiguous contamination in SRB embryo proper and suspensor regions captured by LCM. *(A-C) In situ* localization of SRB G563 *(A),* GA2-oxidase *(B),* and ent-kaurene oxidase (*C*) mRNAs in globular stage seeds. The *G563* gene encodes a protein related to a mung bean *(Vigna radiata)* keratin type 1 cytoskeletal-like protein (BLASTP, 1E-9, 38% amino acid identity). *In situ* hybridization procedures used for SRB seeds have been described in detail elsewhere (1). Images in *(A)* and (*C*) were taken from previously published results by our laboratory under author reuse permissions given by the American Society of Plant Biology (4) and the Proceedings of the National Academy of Sciences (5), respectively. ENT, endothelium. *(D)* Genome browser visualization of SRB G563, GA2-oxidase, and ent-kaurene oxidase RNA-Seq reads in globular stage suspensor (SUS) and embryo proper (EP) regions. Each panel shows the genomic region where the gene is located, including 1 kb of 5’ and 3’ flanking regions. Gene structures and transcription directions (arrows) are shown at the bottom. Numbers indicate RPKM values for each biological replicate. *(E)* RPKM values of the endosperm-specific AGL62 mRNA (6) in SRB suspensor and embryo proper regions.

**Fig. S3.**
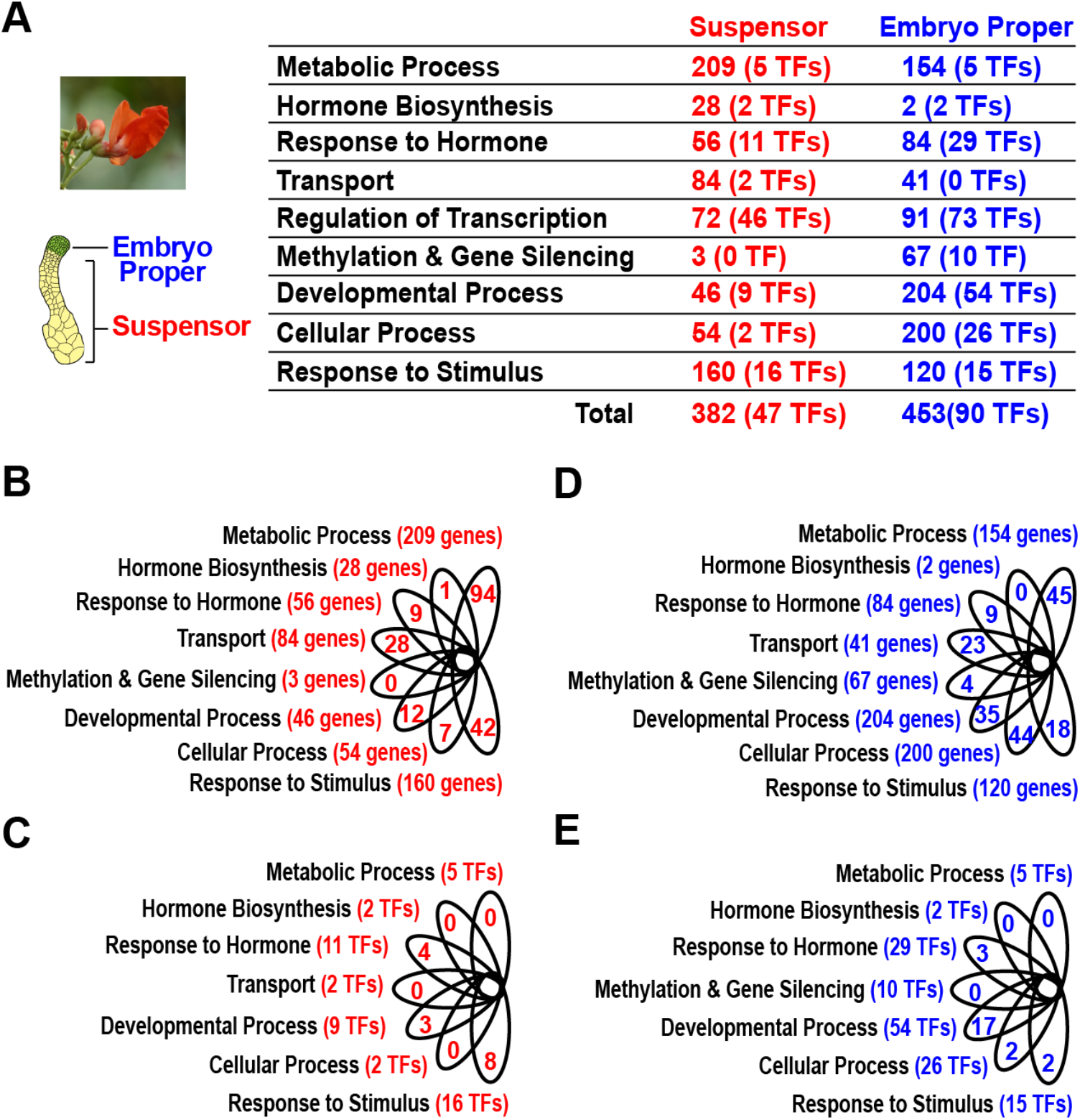
SRB major GO term category up-regulated mRNAs. (*A*) Number of overrepresented up-regulated mRNAs in each GO term category (Fig. 4). (*B-E*) Venn diagrams showing number of up-regulated mRNAs that are unique to each suspensor and embryo proper GO term functional groups. TFs, transcription factors.

**Fig. S4.**
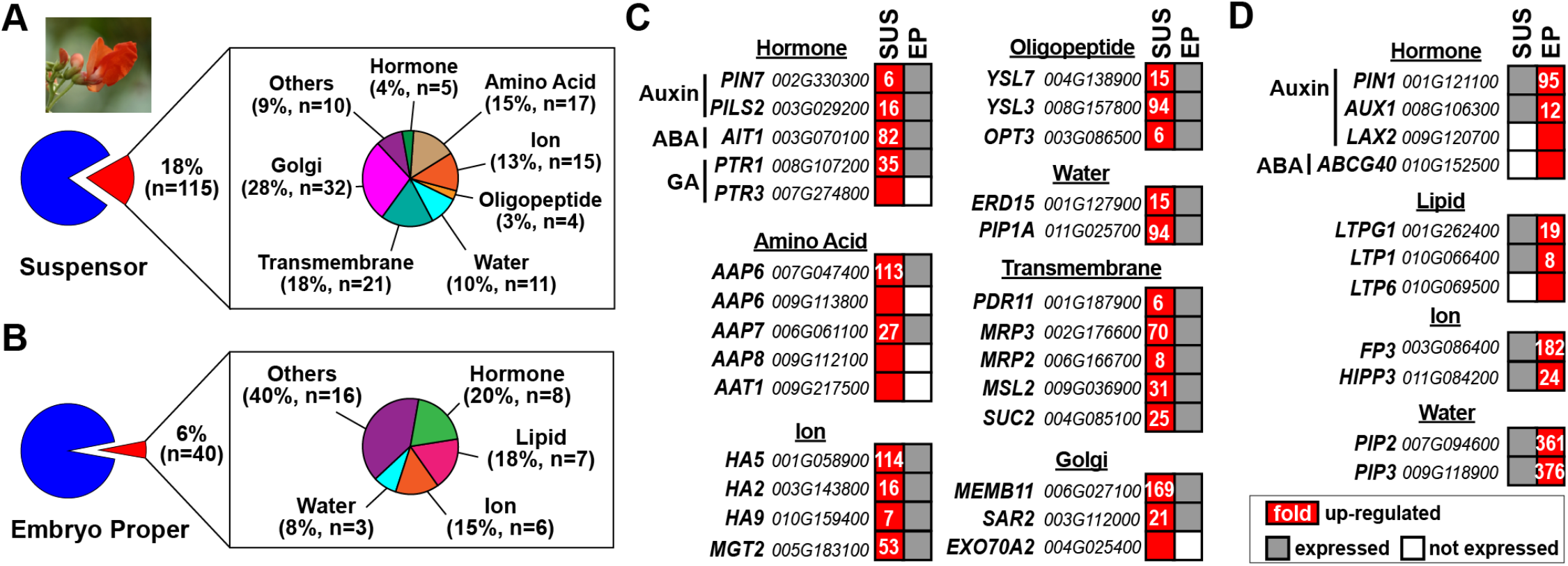
Up-regulated transporter mRNAs in SRB suspensor and embryo proper regions. *(A* and *B*) Percentage of representative transporter mRNAs in suspensor *(A)* and embryo proper (*B*). (*C* and *D*) Up-regulated transporter mRNAs in the suspensor (*C*) and embryo proper (*D*). Numbers in red squares indicate fold differences. See *SI Appendix* Table S1 for gene abbreviations.

**Fig. S5.**
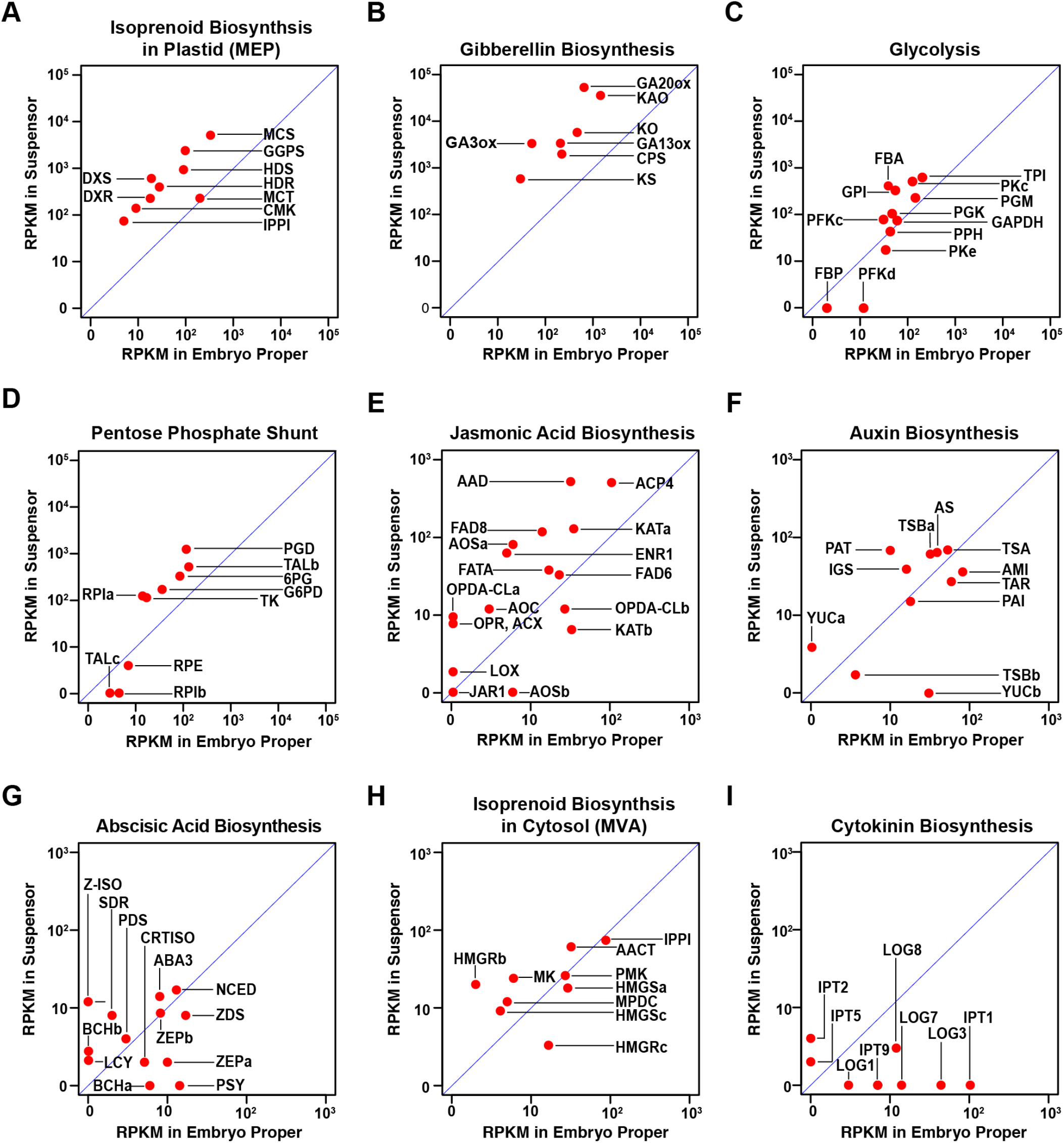
Summary of metabolic pathway enzyme mRNA representation in SRB suspensor and embryo proper regions. Metabolic pathways and enzymes are presented in Fig. 5 and *SI Appendix* Fig. S6, Fig. S7, and Fig. S8. Suspensor and embryo proper RPKM values were plotted for each enzyme mRNA (Dataset S1). Enzyme mRNAs above and below the diagonal line represent those up-regulated in suspensor and embryo proper regions, respectively. Distances from the diagonal line indicate up-regulation levels. The GA biosynthesis mRNAs have the highest levels of suspensor upregulation (Fig. 2 and Fig. 5).

**Fig. S6.**
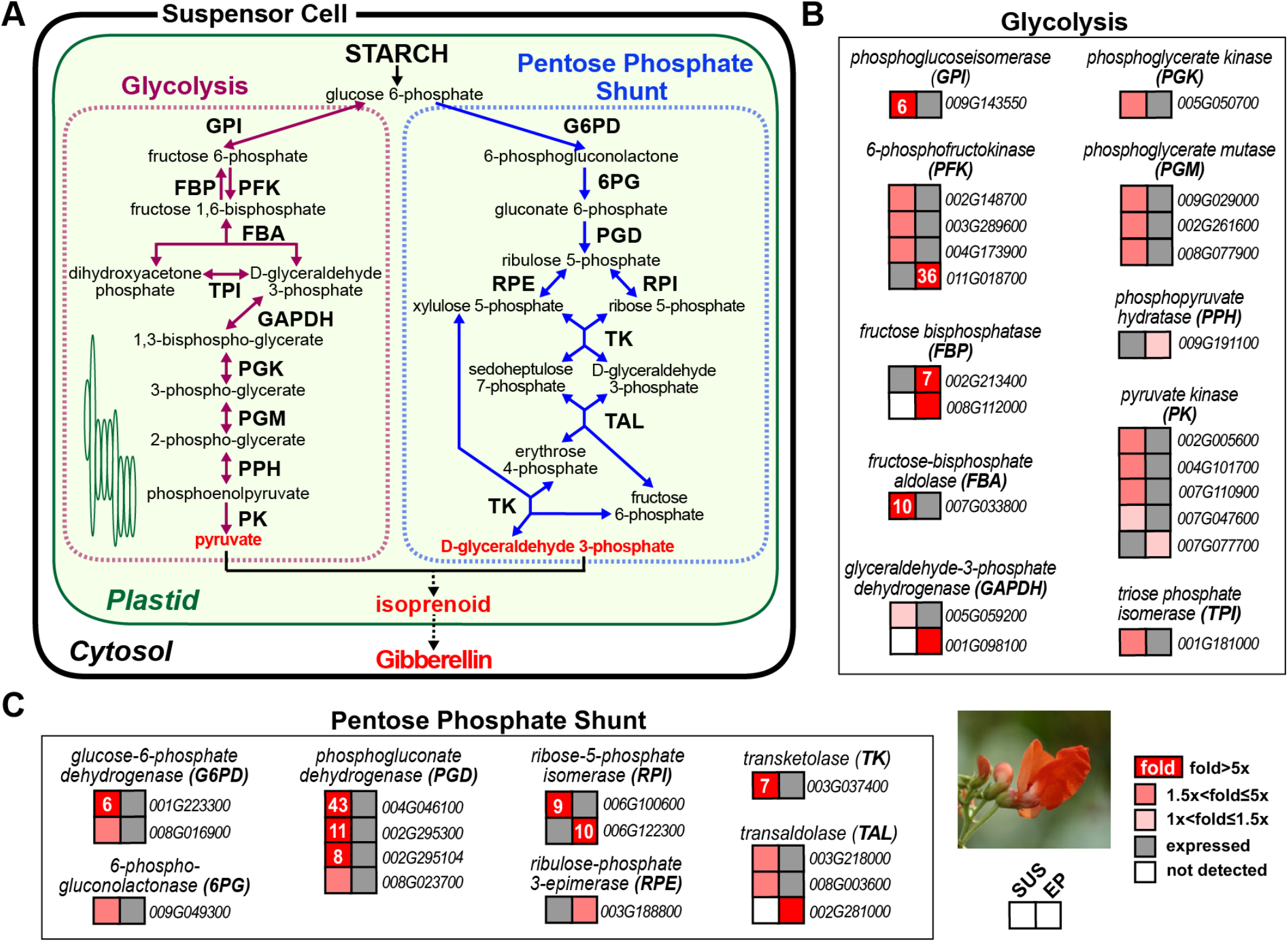
Representation of SRB glycolysis and pentose phosphate shunt enzyme mRNAs in globular stage embryo proper and suspensor regions. (*A*) Glycolysis and pentose phosphate shunt pathways (7–9). The DeepLoc machine learning tool (10) confirmed that enzymes in each pathway were localized in plastids (see Materials and Methods and Dataset S3). (*B* and *C*) Glycolysis (*B*) and pentose phosphate shunt (*C*) enzyme mRNA representation in embryo proper and suspensor regions. Numbers in red squares represent fold differences. Red squares without a number represent mRNAs that were not detected in the other region (white squares). Grey squares indicate mRNAs were detected at levels >0.5 RPKM.

**Fig. S7.**
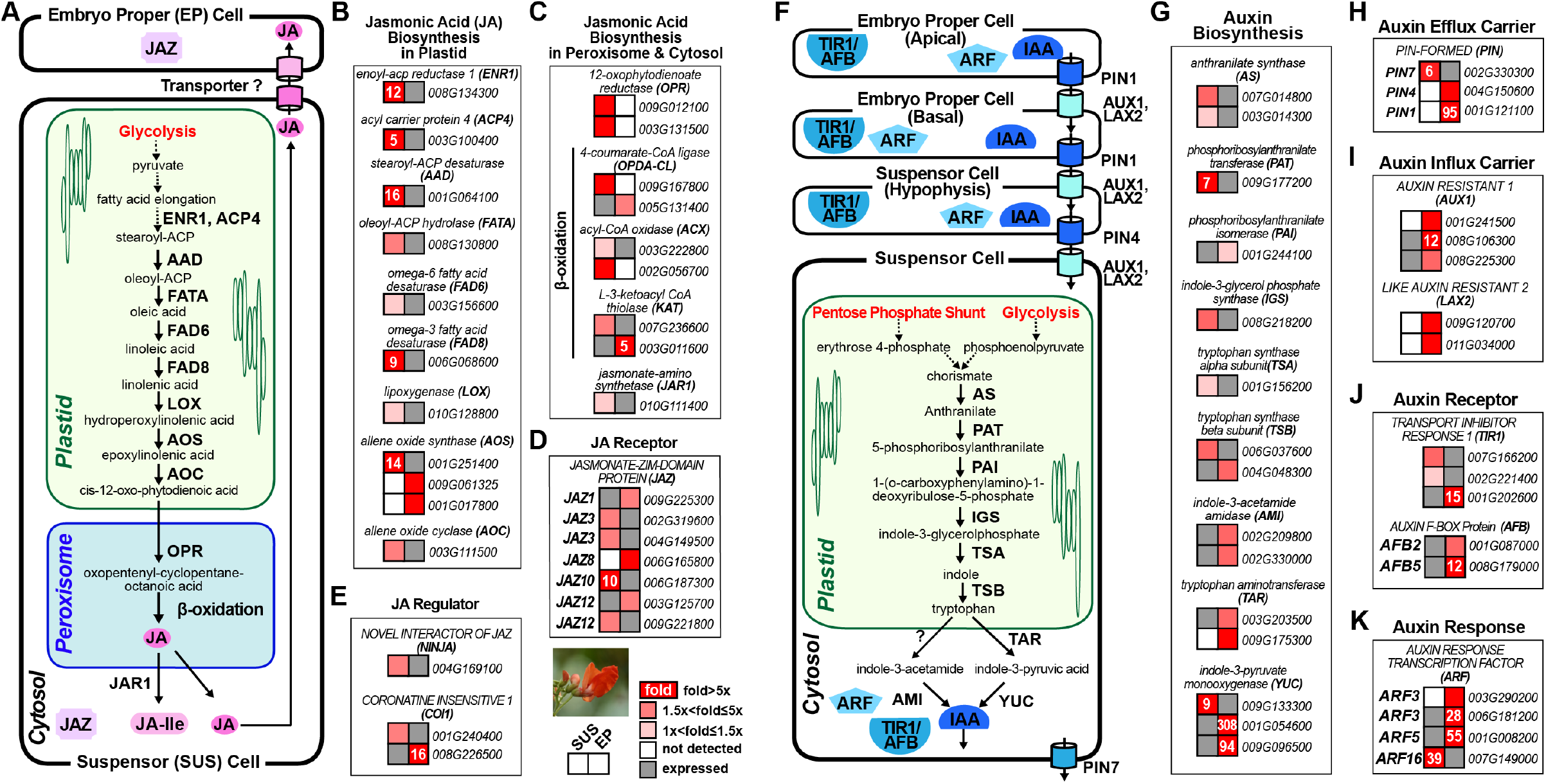
Representation of SRB jasmonic acid (JA) and auxin biosynthesis pathway mRNAs in globular stage embryo proper and suspensor regions. (*A*) JA biosynthesis and signaling pathways (11, 12). Localization of enzymes and proteins in different cellular compartments were confirmed by using the DeepLoc machine learning tool (10) (see Materials and Methods and Dataset S3). (*B-D*) Suspensor and embryo proper mRNAs encoding JA biosynthesis enzymes *(B* and *C*), receptors (*D*), and regulators (*E*). *(F)* Auxin biosynthesis pathway (12, 13). Cellular localizations were confirmed by the DeepLoc learning tool (10) (see Materials and Methods and Dataset S3). Model for auxin metabolic and signal transduction pathways was based on: (i) presence of corresponding mRNAs (*G-K*); (ii) classical data showing that auxin is present in SRB suspensor and embryo proper regions (14); (iii) early hypotheses suggesting auxin is synthesized in the SRB suspensor and transported to the embryo proper (14); and (iv) detailed knowledge of auxin signal transduction pathway in *Arabidopsis* embryos (15). (*G-K*) Suspensor and embryo proper mRNAs encoding auxin biosynthetic enzymes (*G*), efflux carriers (*H*), influx carriers (*I*), receptors (*J*), and response transcription factors (*K*). Numbers in red squares represent fold differences. Red squares without a number represent mRNAs that were not detected in the other region (white squares). Grey squares indicate mRNAs were detected at levels >0.5 RPKM.

**Fig. S8.**
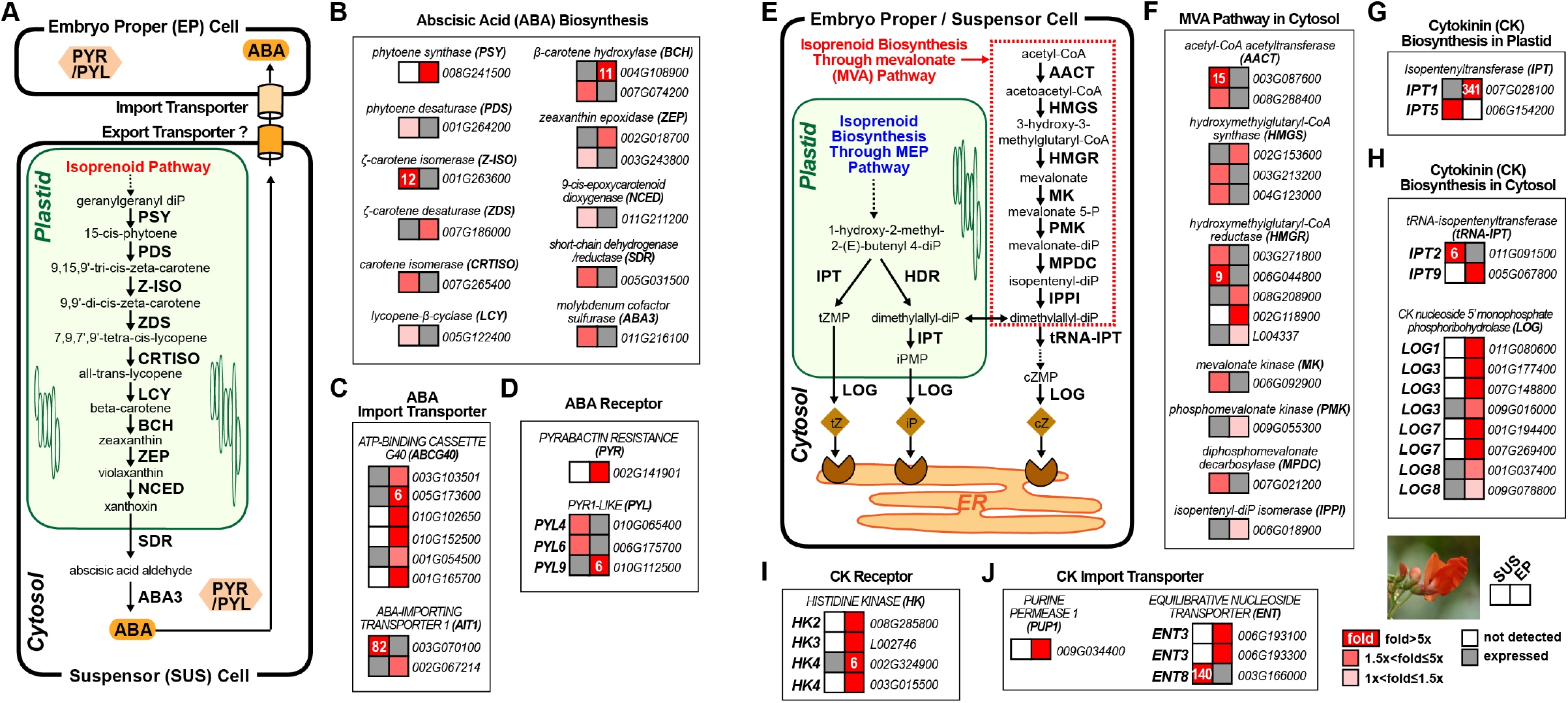
Representation of SRB abscisic acid (ABA) and cytokinin (CK) biosynthesis pathway mRNAs in globular stage embryo proper and suspensor regions. (*A*) ABA biosynthetic and signaling pathways (12). Localization of enzymes and proteins in specific cellular compartments were confirmed by using the DeepLoc machine learning tool (10) (see Materials and Methods and Dataset S3). (*B-D*). Suspensor and embryo proper mRNAs encoding ABA biosynthesis enzymes (*B*), transporters (*C*), and receptors (*D*). (*E*) CK biosynthetic pathway (12). Localization of enzymes and proteins in specific cellular compartments were confirmed by using the DeepLoc machine learning tool (10) (see Materials and Methods and Dataset S3). (*F-J*). Suspensor and embryo proper mRNAs encoding CK biosynthesis enzymes (*F-H*), receptors (*I*), and transporters (*J*). Numbers in red squares represent fold differences. Red squares without a number represent mRNAs that were not detected in the other region (white squares). Grey squares indicate mRNAs were detected at levels >0.5 RPKM. tZ, *trans-Zeatin;* cZ, *cis*-Zeatin; iP, *N^6^*-(^2^-Isopentenyl) adenine.

**Fig. S9.**
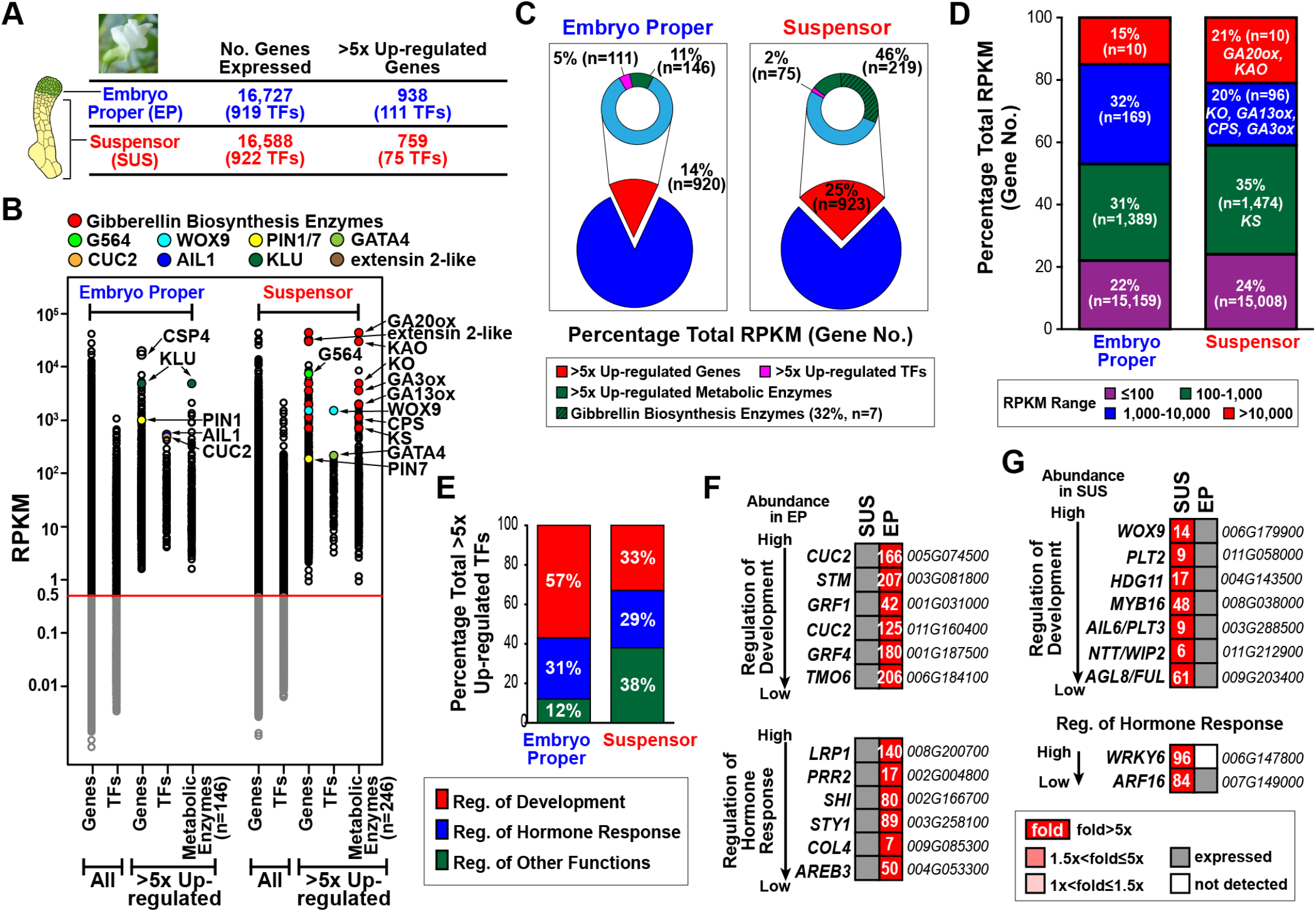
Gene activity in CB suspensor and embryo proper regions. *(A)* Genes active (>0.5 RPKM) in at least one biological replicate, and >5 fold up-regulated genes in embryo proper and suspensor (see Materials and Methods). All expressed genes and those up-regulated >5-fold are listed in Dataset S1. TFs, transcription factor mRNAs. *(B)* Abundance distribution of mRNAs in different embryo proper and suspensor mRNA populations represented by RPKM. Each circle represents a different mRNA sequence. RNAs with <0.5 RPKM in both biological replicates were not used in subsequent analyses, as indicated by the grey circles below the red line. Enzyme mRNAs were identified using the Plant Metabolic Pathway Database (https://www.plantcyc.org) (Dataset S3). (*C*) Percentage of total RPKM for different embryo proper and suspensor mRNA populations. Parentheses represent the number of mRNAs in each population. *(D)* Percentage of embryo proper and suspensor mRNA populations with different prevalence levels. Parentheses indicate the number of different mRNAs in each prevalence group. *(E)* Percentage of TF mRNAs in different functional groups *(SI Appendix* Fig. S10). *(F* and *G)* Representative up-regulated embryo proper (*F*) and suspensor (*G*) TF mRNAs listed by prevalence levels. Numbers in red squares represent fold differences. Red squares without a number indicate that mRNAs were not detected in the other embryo region (white squares). Grey squares indicate mRNAs were detected at levels >0.5 RPKM. Gene identifiers (i.e., Pv number) given to right. Gene abbreviations are defined in *SI Appendix* Table S1.

**Fig. S10.**
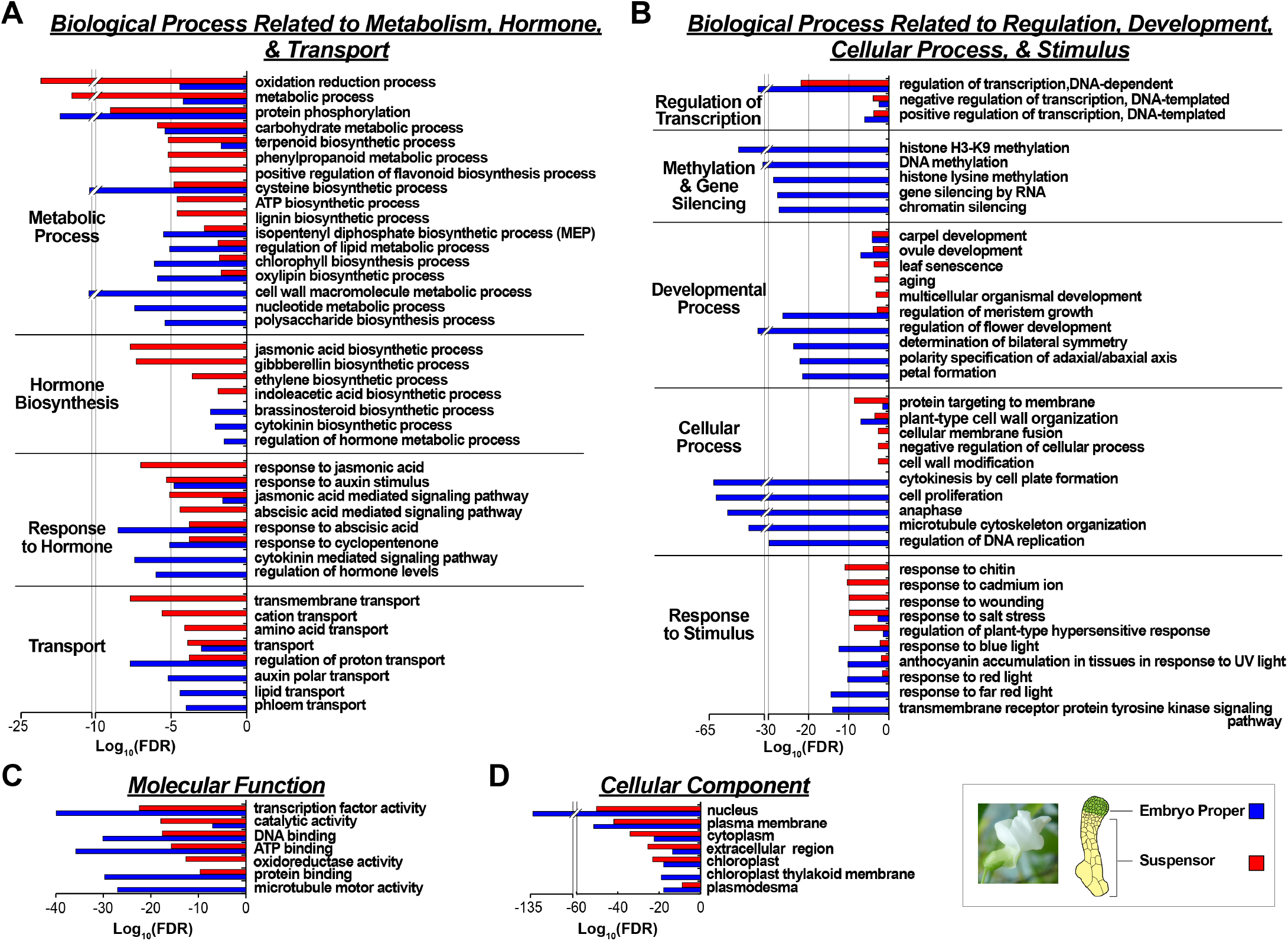
GO terms that are enriched in CB suspensor and embryo proper regions. *(A* and *B)* Enriched biological process GO terms related to metabolism, hormone, and transport *(A)* and regulation, development, cellular process, and stimulus (*B*). *(C* and *D)* Molecular function (*C*), and cellular component (*D*) GO terms. Only the top five to 10 GO terms are listed. All GO terms are presented in an interactive format in Dataset S2. FDR, false discovery rate.

**Fig. S11.**
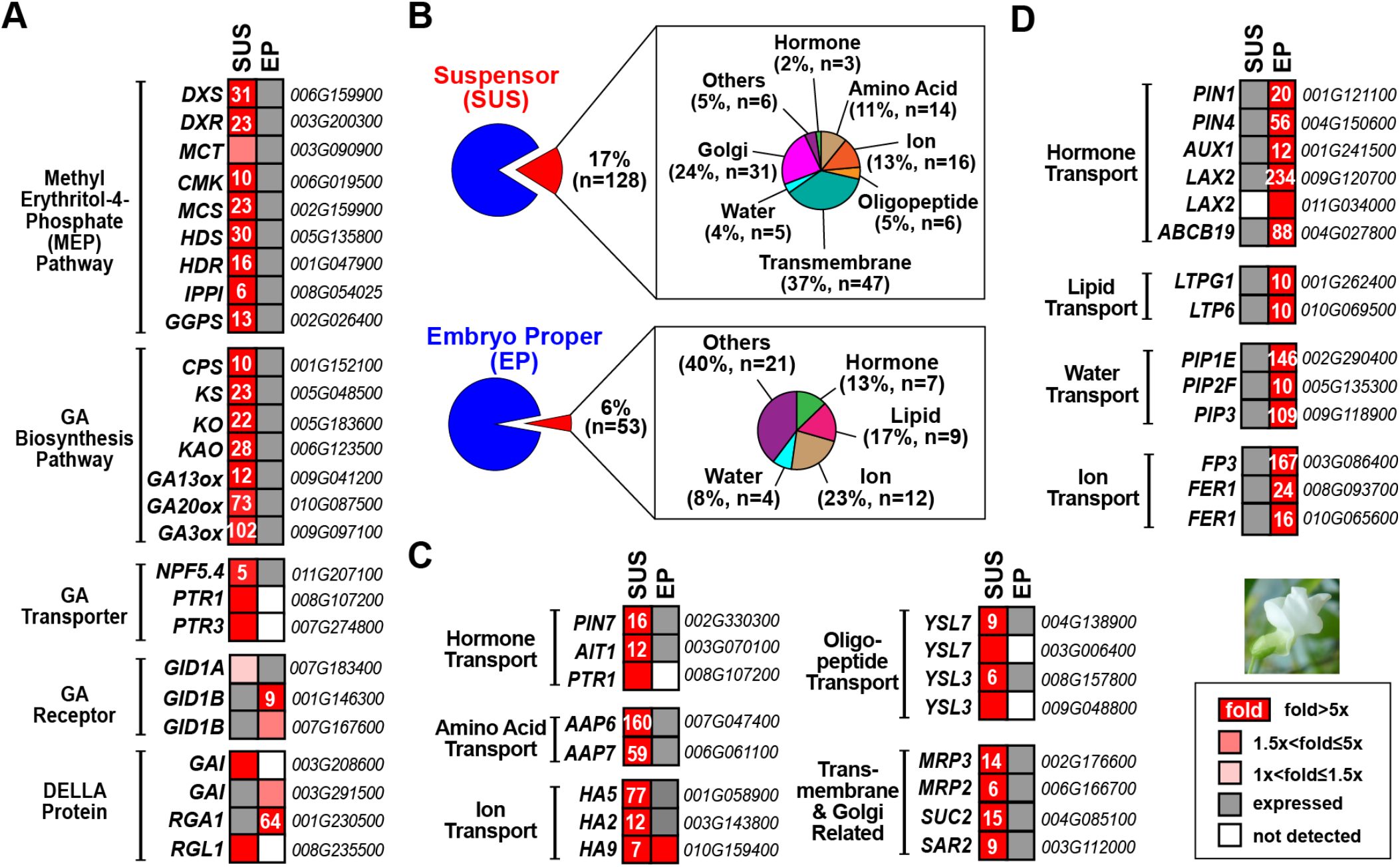
Representation of CB MEP, GA biosynthesis, and transporter mRNAs in embryo proper and suspensor regions. (*A*) MEP and GA biosynthesis pathway and signaling mRNAs. (*B*) Percentage of representative transporter mRNAs in suspensor and embryo proper regions. (*C* and *D*) Up-regulated transporter mRNAs in the suspensor (*C*) and embryo proper (*D*). Numbers in red squares represent fold differences. Red squares without a number indicate that mRNAs were not detected in the other embryo region (white squares). Grey squares indicate mRNAs were detected at levels >0.5 RPKM. Gene identifiers (i.e., Pv number) given to right. Gene abbreviations are defined in *SI Appendix* Table S1.

**Fig. S12.**
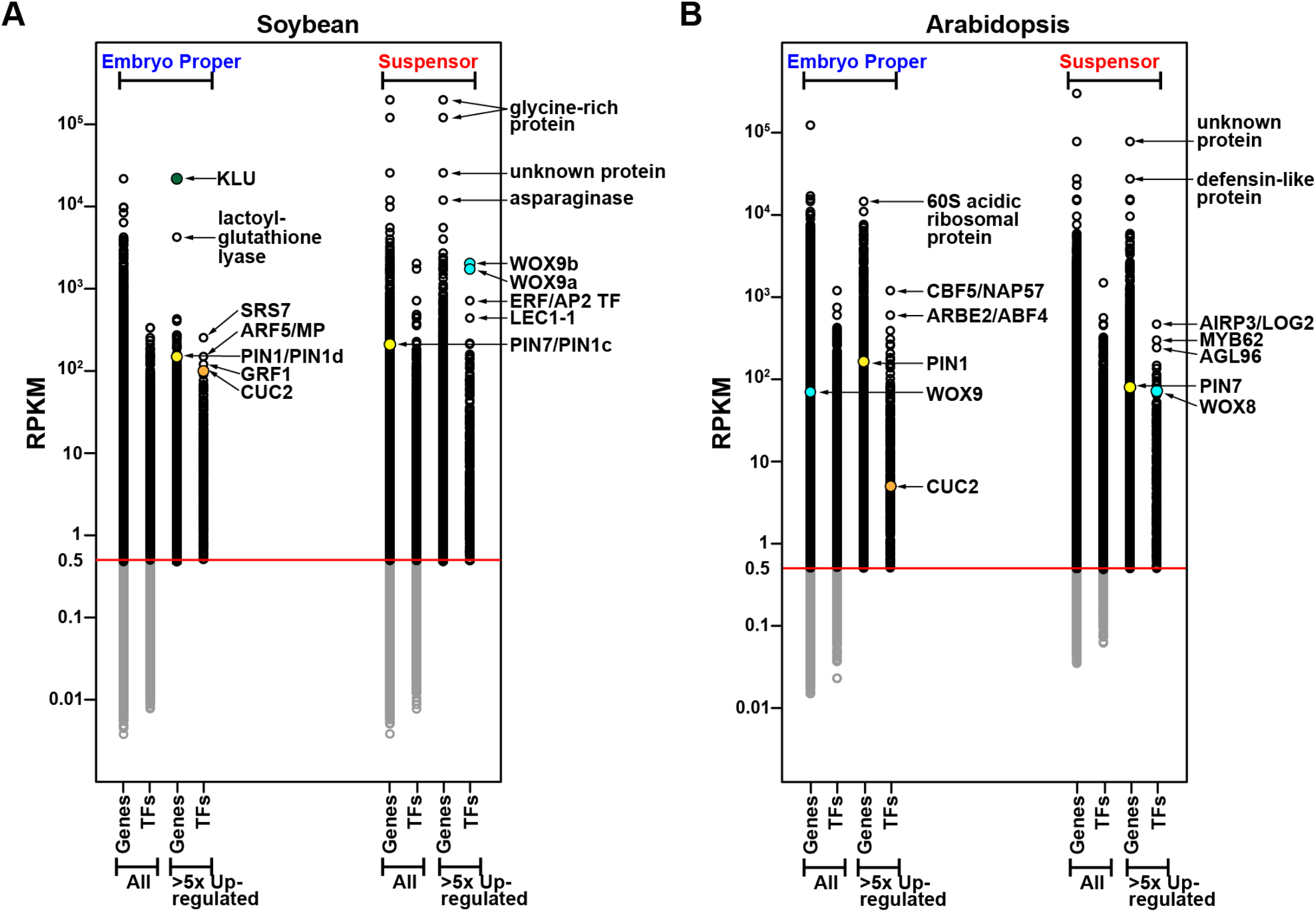
Abundance distribution of SB and *Arabidopsis* embryo mRNAs. (*A* and *B*) Abundance distribution of SB (*A*) and *Arabidopsis* (*B*) mRNAs in embryo proper and suspensor mRNA populations represented by RPKM (Dataset S1). Each circle represents a different mRNA sequence. RNAs with <0.5 RPKM in both biological replicates were not used in subsequent analyses, as indicated by the grey circles below the red line. PIN1d and PIN1c refer to most closely related SB *PIN* gene family members (16). We designated these as PIN1 and PIN7 because they have embryo proper and suspensor localization patterns similar to *Arabidopsis* PIN1 and PIN7 mRNAs (17).

**Fig. S13.**
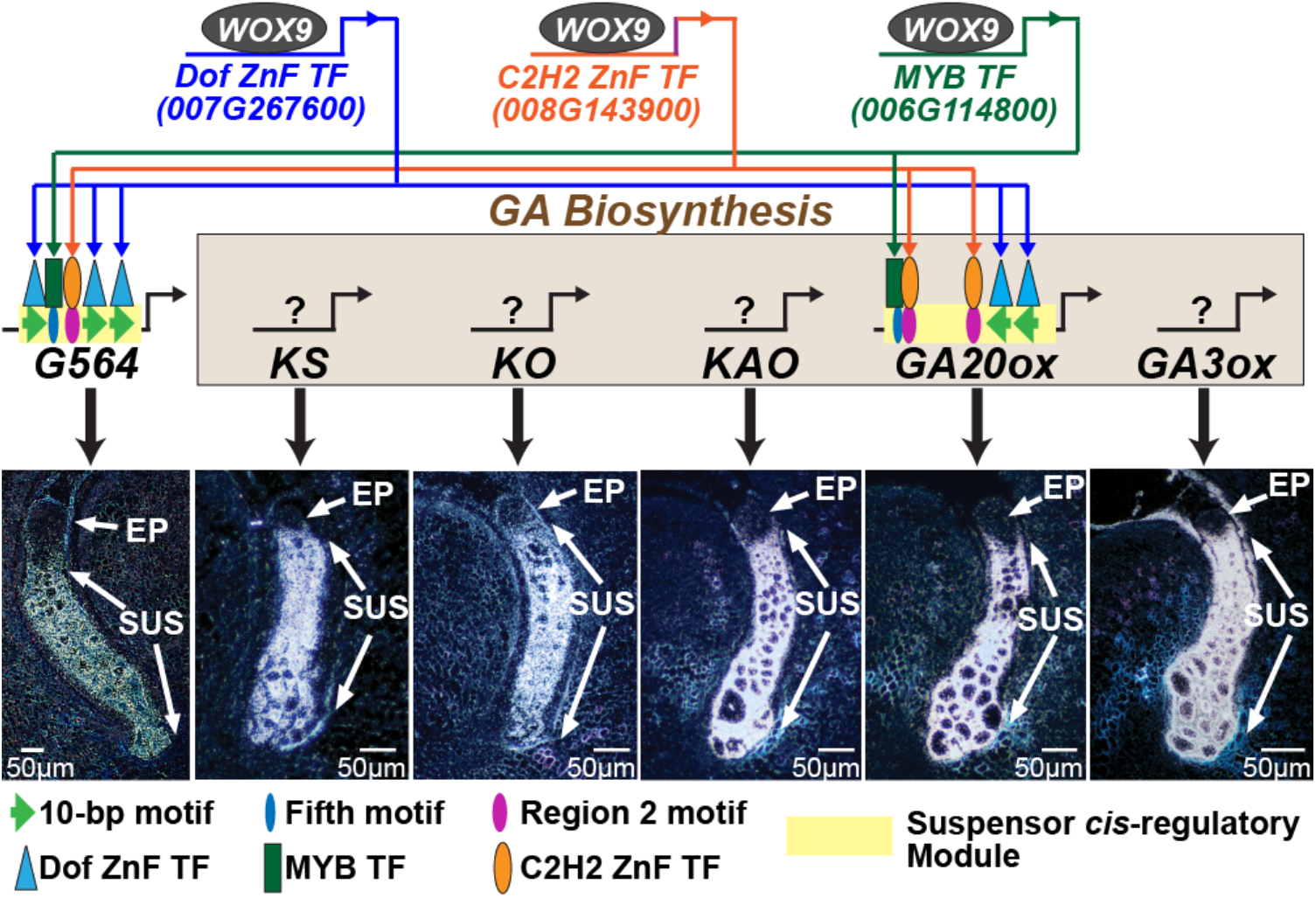
Model for *GA20ox* and *G564* gene activation within the suspensor. Model is based on the WOX9 ChIP-Seq experiments reported here (Fig. 9) and previous experiments in our laboratory on *GA20ox* and *G564* suspensor *cis* regulatory modules (5, 18). G564 and GA enzyme mRNA localization images in the SRB suspensor were taken from previously published results by our laboratory under author reuse permissions given by the American Society of Plant Biology (G564) (4) and the Proceedings of the National Academy of Sciences (GA enzymes) (5). ZnF, zinc finger.

**Table S1.**
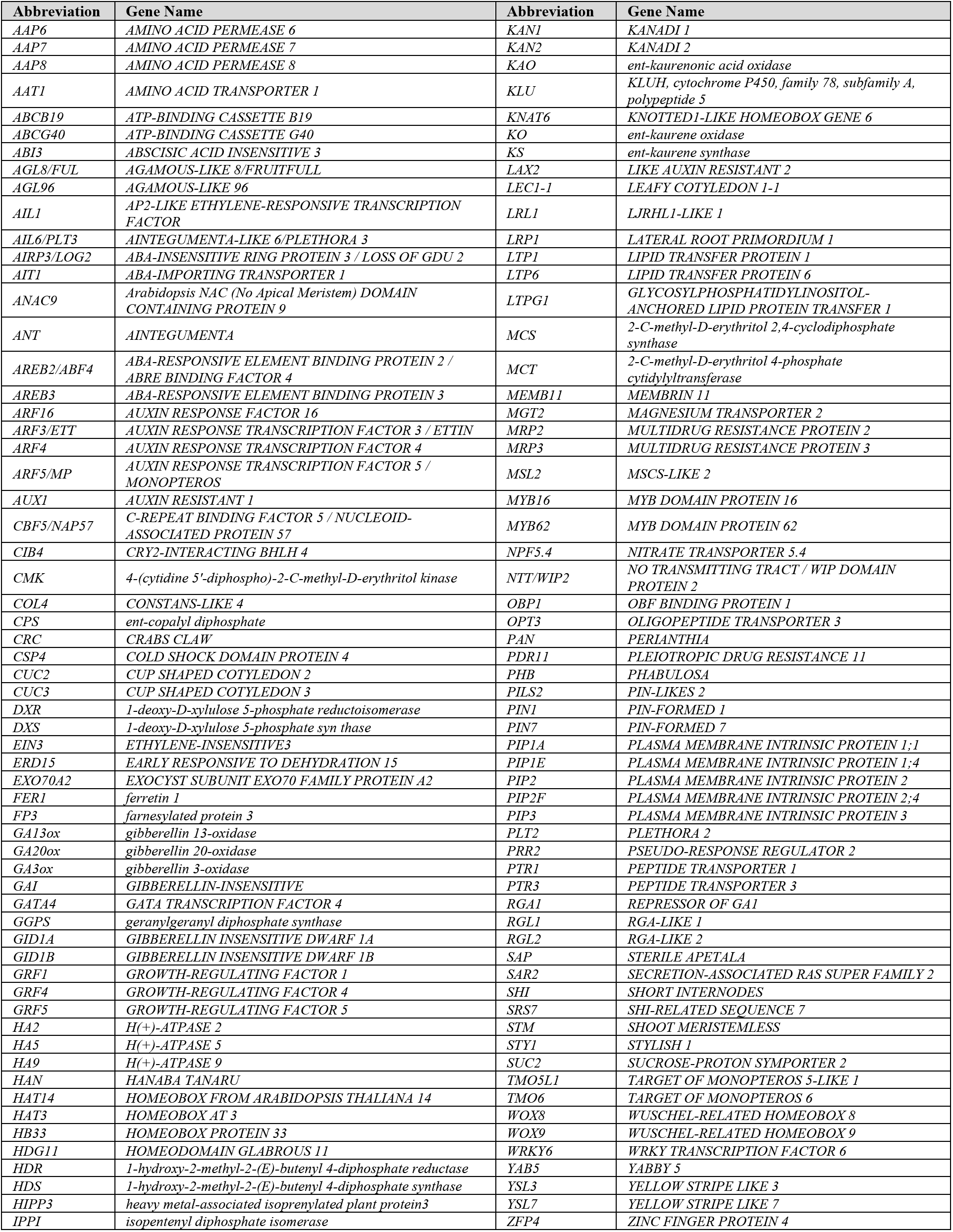
Gene Abbreviations in Figures.

**Table S2.**
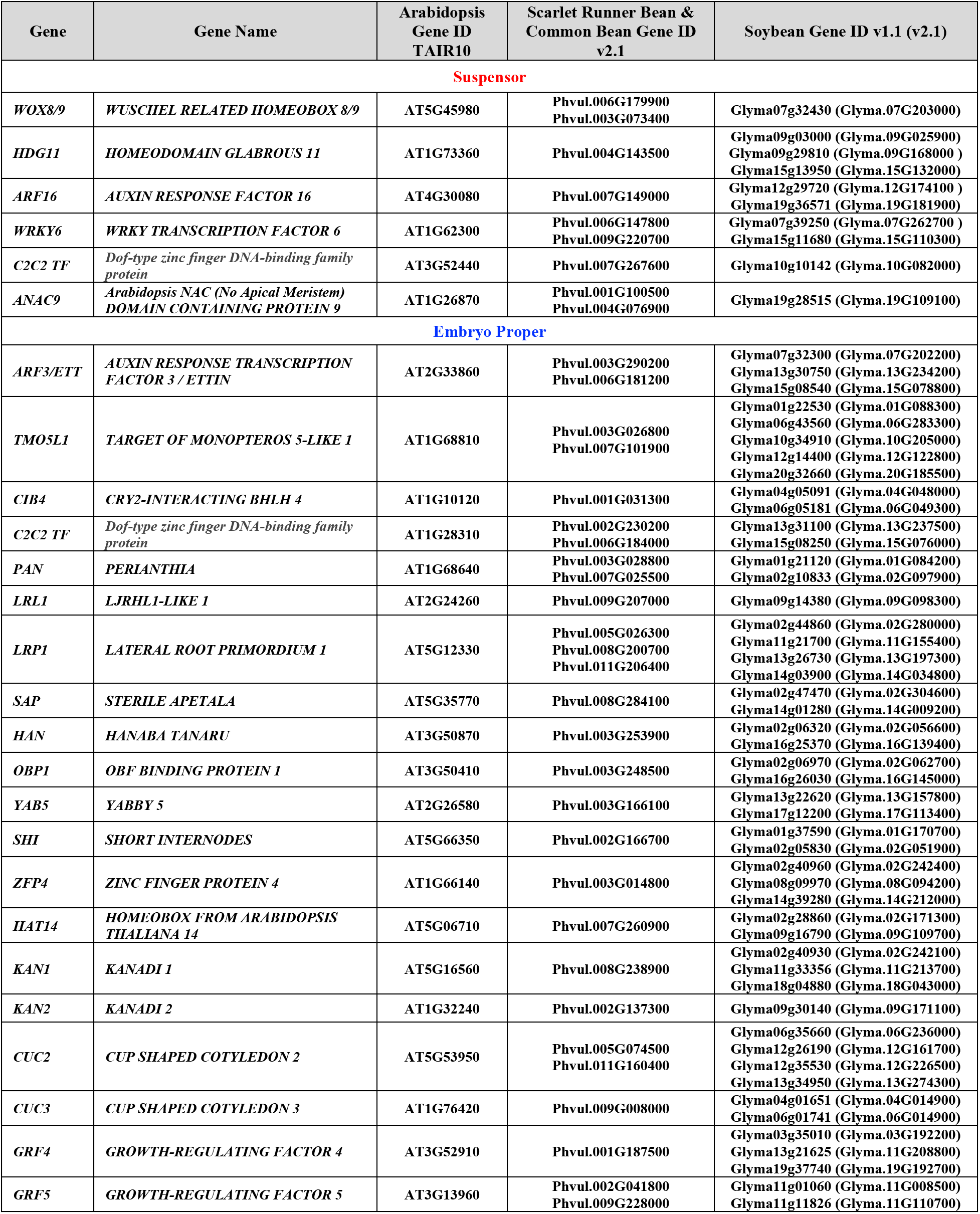
Shared Up-Regulated Transcription Factor mRNAs in Suspensor and Embryo Proper Regions.

**Dataset S1.**
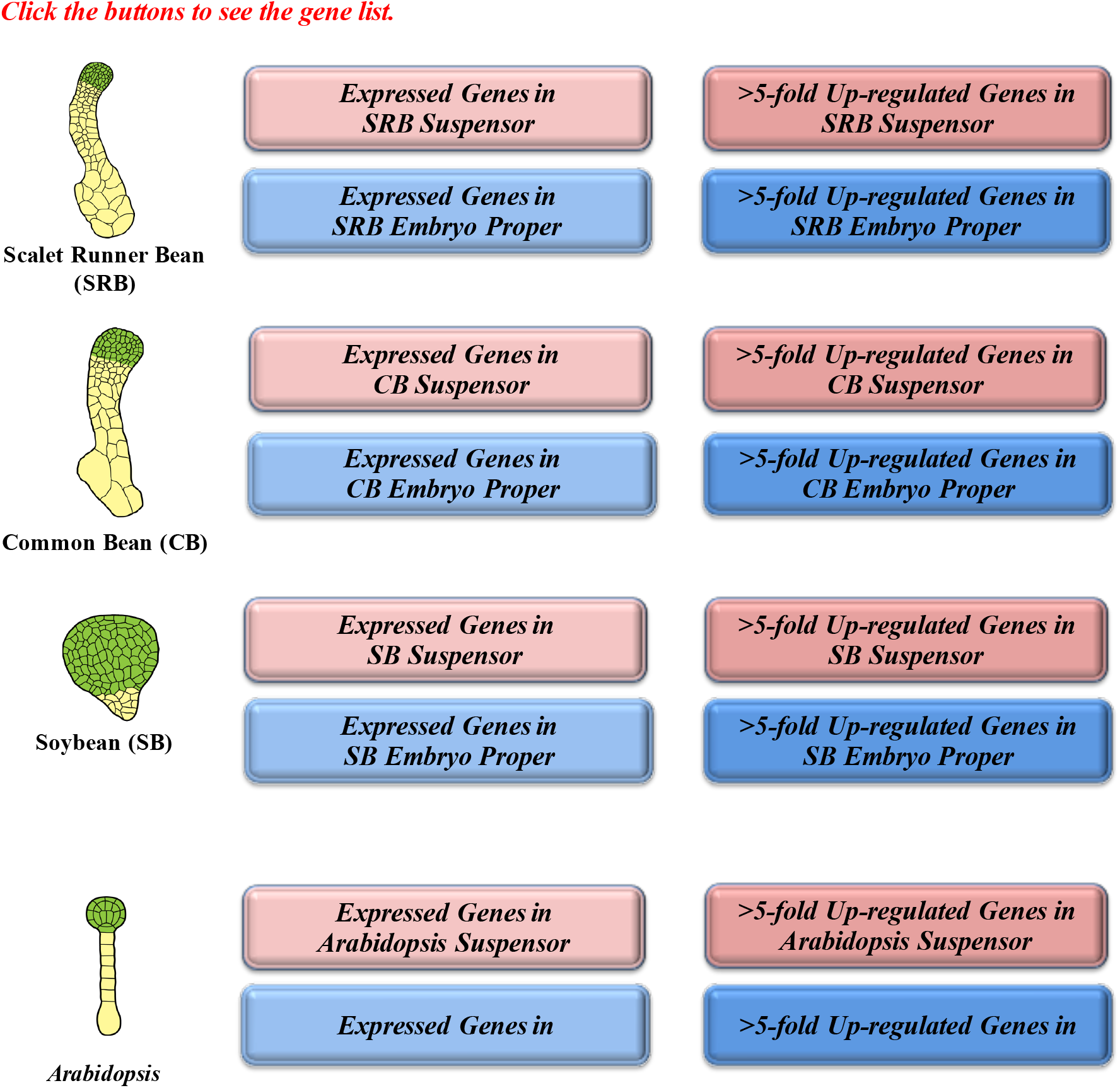
Expressed and >5-fold Up-regulated Genes* in Scarlet Runner Bean (SRB), Common Bean (CB), Soybean (SB), and Arabidopsis Globular Stage Suspensor and Embryo Proper * Up-regulated genes were identified by edgeR package with FDR<0.05 and >5-fold change.

**Dataset S2.**
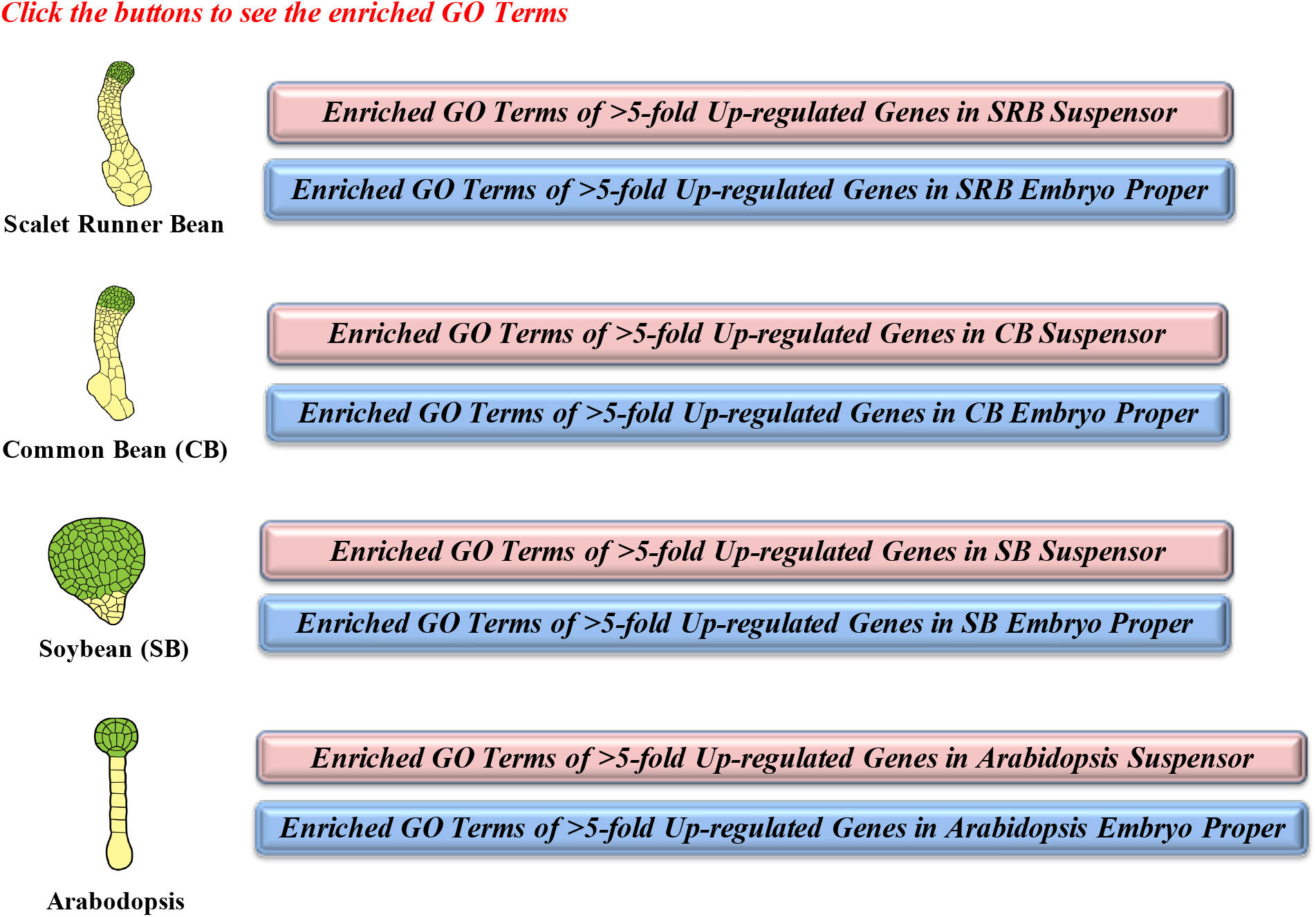
Enriched GO Terms of >5-fold up-regulated genes in suspensor and embryo proper of Scarlet Runner Bean, Common Bean, Soybean, and Arabidopsis

**Dataset S3.**
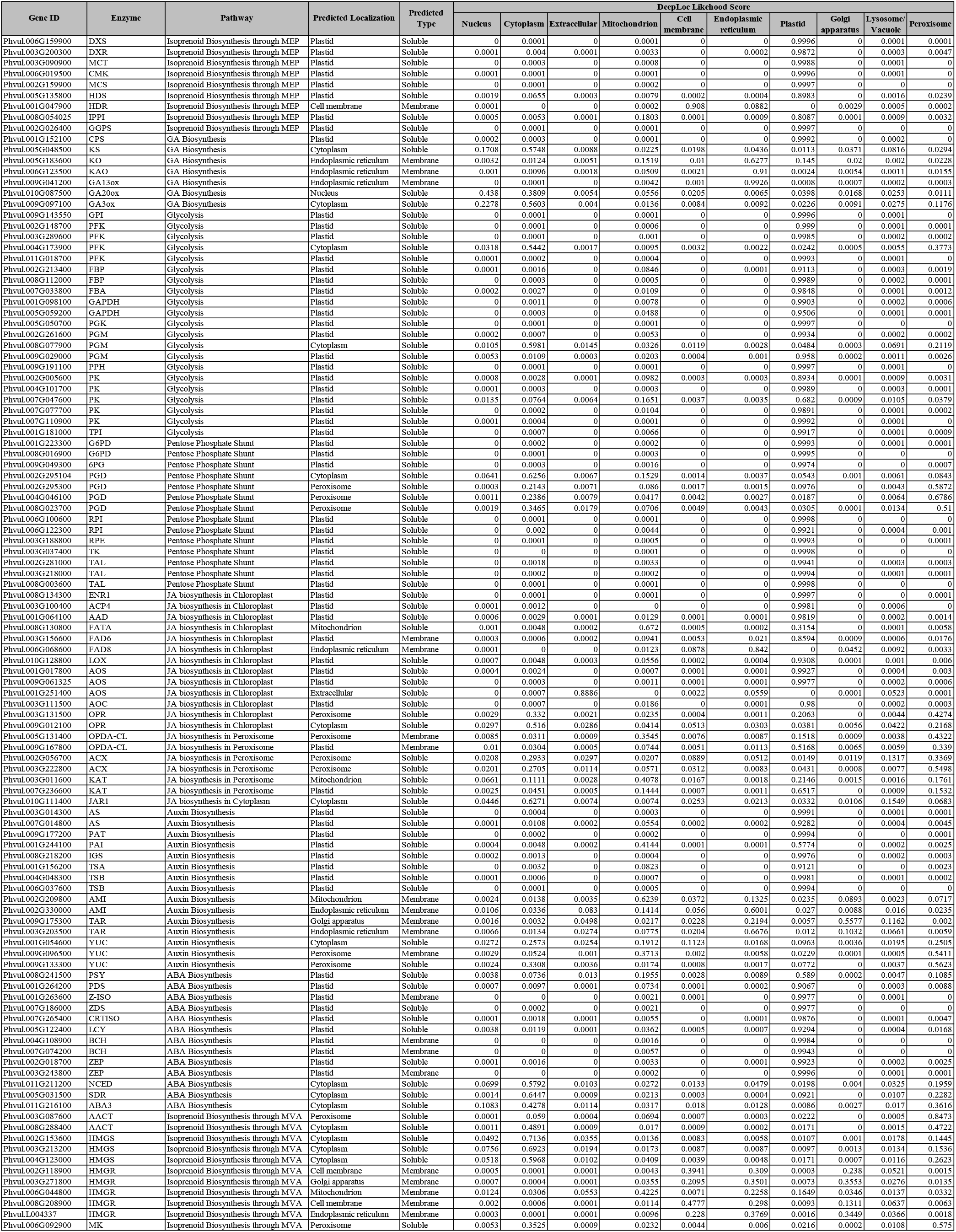
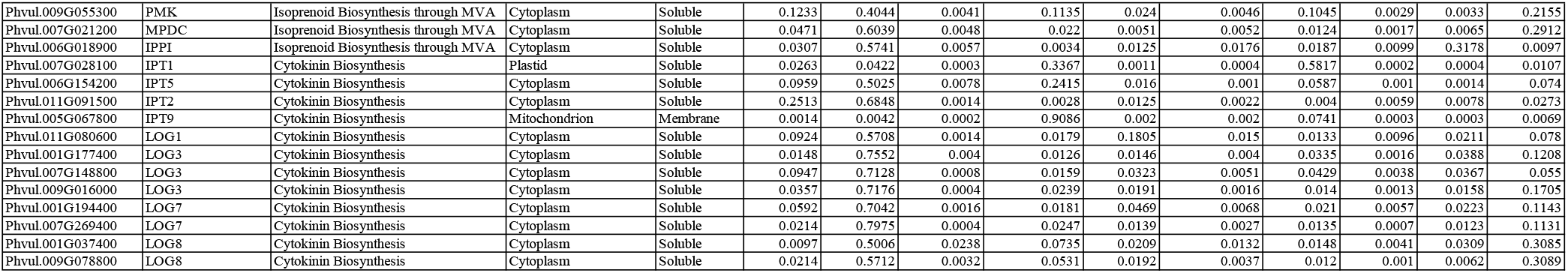
Subcellular Localization Prediction of Enzymes in Metabolic Pathways in Scarlet Runner Bean and Common Bean. The Plant Metabolic Pathway Database (v13) (https://www.plantcyc.org) was used to identify metabolic pathways encoded by up-regulated suspensor and embryo proper enzyme mRNAs. Enzyme protein sequences were used to predict their subcellular locations using the machine learning program DeepLoc-1.0 (http://www.cbs.dtu.dk/services/DeepLoc/).

**Dataset S4.**
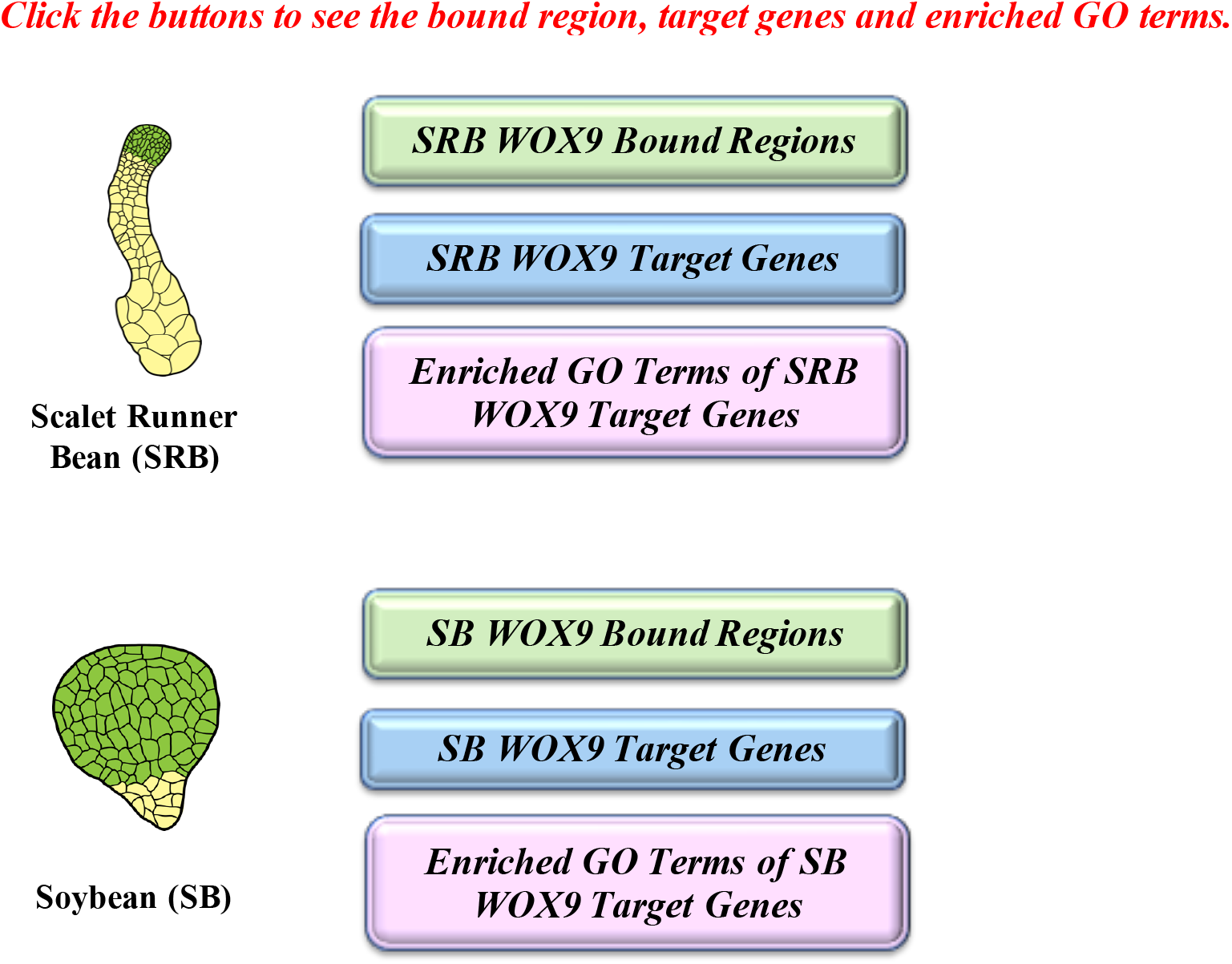
Scarlet Runner Bean and Soybean WOX9 Bound Regions and Target Genes.

